# The influence of peptide context on signalling and trafficking of glucagon-like peptide-1 receptor biased agonists

**DOI:** 10.1101/2020.02.24.961524

**Authors:** Zijian Fang, Shiqian Chen, Philip Pickford, Johannes Broichhagen, David J Hodson, Ivan R Corrêa, Sunil Kumar, Frederik Görlitz, Christopher Dunsby, Paul French, Guy A Rutter, Tricia Tan, Stephen R Bloom, Alejandra Tomas, Ben Jones

## Abstract

Signal bias and membrane trafficking have recently emerged as important considerations in the therapeutic targeting of the glucagon-like peptide-1 receptor (GLP-1R) in type 2 diabetes and obesity. In the present study, we have evaluated a peptide series with varying sequence homology between native GLP-1 and exendin-4, the archetypal ligands on which approved GLP-1R agonists are based. We find notable differences in agonist-mediated signalling, endocytosis and recycling, dependent both on the introduction of a His → Phe switch at position 1 and the specific mid-peptide helical regions and C-termini of the two agonists. These observations were linked to insulin secretion in a beta cell model and provide insights into how ligand factors influence GLP-1R function at the cellular level.

**Graphical abstract:** 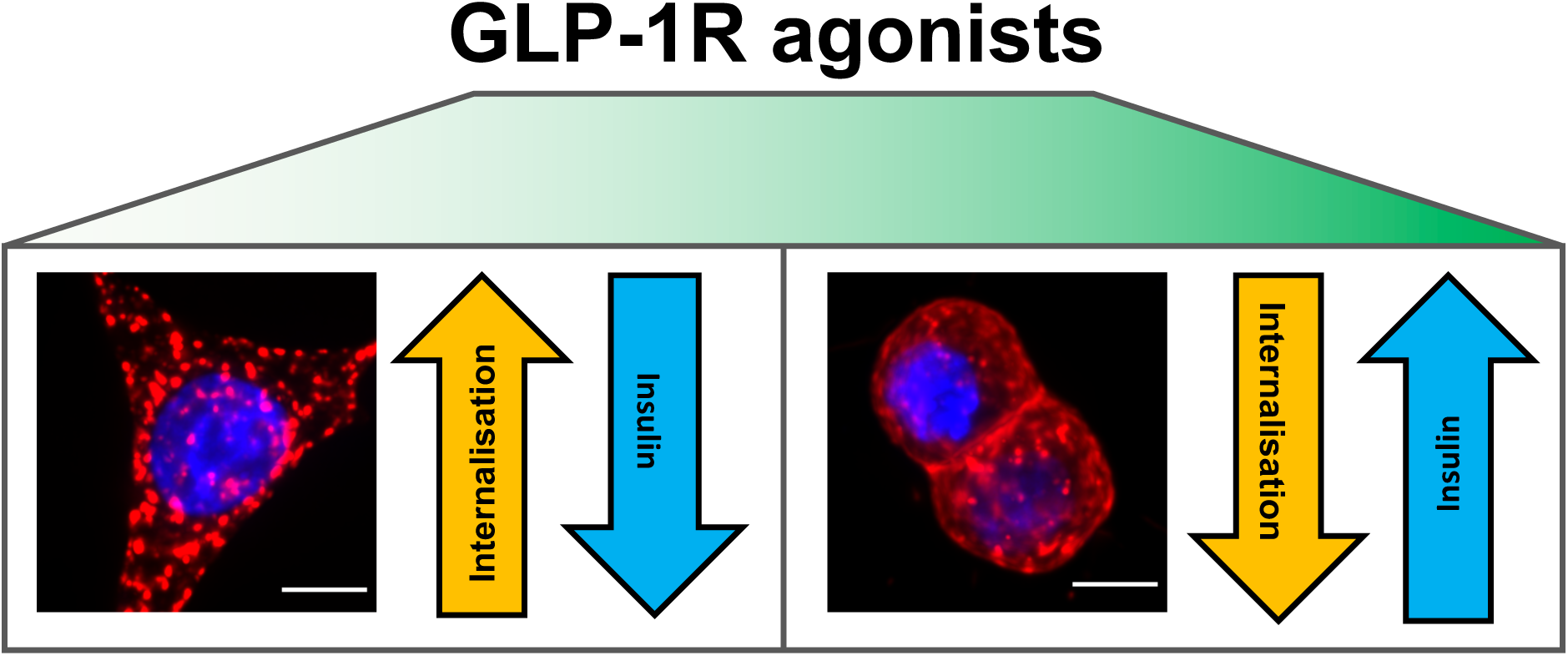

## 1 Introduction

Due to the increasing worldwide prevalence of both type 2 diabetes (T2D) and obesity (1, 2), there is considerable interest in the identification and optimisation of drugs which can treat both of these conditions. The glucagon-like peptide-1 receptor (GLP-1R) is expressed in pancreatic beta cells and anorectic neurons in the brain, and promotes insulin secretion and weight loss when activated by endogenous or therapeutic peptide ligands (3). Consequently, GLP-1R agonists (GLP-1RAs) are commonly used to treat T2D and related metabolic diseases (4).

Activated GLP-1Rs engage with cytosolic effectors to generate intracellular signalling responses such as production of cyclic adenosine monophosphate (cAMP), elevations in intracellular calcium (Ca^2+^), recruitment of β-arrestins and phosphorylation of extracellular regulated kinase (ERK) (5). Ligand-specific signalling pathway preference (“signal bias”) has emerged as a factor controlling downstream GLP-1R actions such as potentiation of insulin secretion (6), and is of ongoing interest in the therapeutic targeting of other membrane receptors as it provides a potential means to accentuate desirable effects and minimise side effects (7). Moreover, endocytosis and post-endocytic trafficking influence the availability of GLP-1Rs at the cell surface and fine-tune the spatiotemporal origin of signalling responses (8–10).

The GLP-1 homologue peptide exendin-4 was the first therapeutic GLP-1RA developed for clinical use (11). Exendin-4 is a high affinity agonist with enhanced resistance to proteolytic degradation in comparison to native GLP-1 (12). Three recent studies indicate how N-terminal amino acid sequence changes to exendin-4 can improve its metabolic effects by generating signalling response profiles that accentuate cAMP generation over β-arrestin recruitment and/or GLP-1R internalisation (13–15). However, it is not known if similar effects can be achieved by modifying the N-terminus of native GLP-1. As the amino acid sequences of a number of approved GLP-1RAs are highly similar to GLP-1 itself, e.g. Liraglutide, Dulaglutide and Semaglutide (16), this question is of potential therapeutic importance. Furthermore, recent data suggests a potential advantage for GLP-1-like agents over exendin-4-based agonists for clinical outcomes, raising the possibility that the pharmacology of these two agonist sub-classes is intrinsically non-identical (17, 18). A deeper understanding of GLP-1RA structural features that control signal bias and trafficking may aid in the development of better drugs to treat T2D.

In this report we have tested a panel of chimeric peptide GLP-1RAs carrying features of both native GLP-1 and exendin-4, as well as their derivatives modified with a His → Phe switch at position 1. In the context of exendin-4, the latter single amino acid change was previously shown to result in favourable pharmacological characteristics including faster dissociation kinetics, reduced β-arrestin recruitment and endocytosis, faster GLP-1R recycling, and greater insulin secretion *in vitro* and *in vivo* (14). Here, we use a variety of *in vitro* approaches to demonstrate that peptide-triggered receptor signalling and trafficking properties are influenced by relative homology to GLP-1 *versus* exendin-4, with introduction of exendin-4-specific mid-peptide helical sequences associated with slower GLP-1R recycling and greater desensitisation. The effects of the His1 → Phe1 switch are also modulated by contextual sequence differences. These findings highlight the importance of the entire peptide sequence in the development of improved biased ligands targeting GLP-1R.

## 2 Results

### 2.1 Peptide regions contributing to binding, signalling activity and trafficking responses of exendin-4 and GLP-1 chimeric peptides

We first generated a panel of chimeric peptides bearing features of both GLP-1(7–36)NH_2_ (henceforth referred to as “GLP-1”) and exendin-4 (Table 1, Supplementary Figure 1A). These were modelled on a series described in an earlier report (19). In the latter study, the chimeric peptide N-termini were truncated, in order to discern the relative contribution of different structural features to binding affinity; in the present study the N-termini are intact, albeit with amino acid substitutions compared to the parent peptide in some cases. Chimera 1 (Chi1) contains the full sequence of GLP-1 with the addition of the exendin-4 C-terminus; chimera 2 and 3 (Chi2, Chi3) progressively incorporate more of the exendin-4 sequence within the mid-peptide helical region. Additionally, to probe the role of the penultimate residues of exendin-4 (Gly) and GLP-1 (Ala), these were switched in two peptides to produce Ex-ala2 and GLP-1-gly2.

**Table 1.**
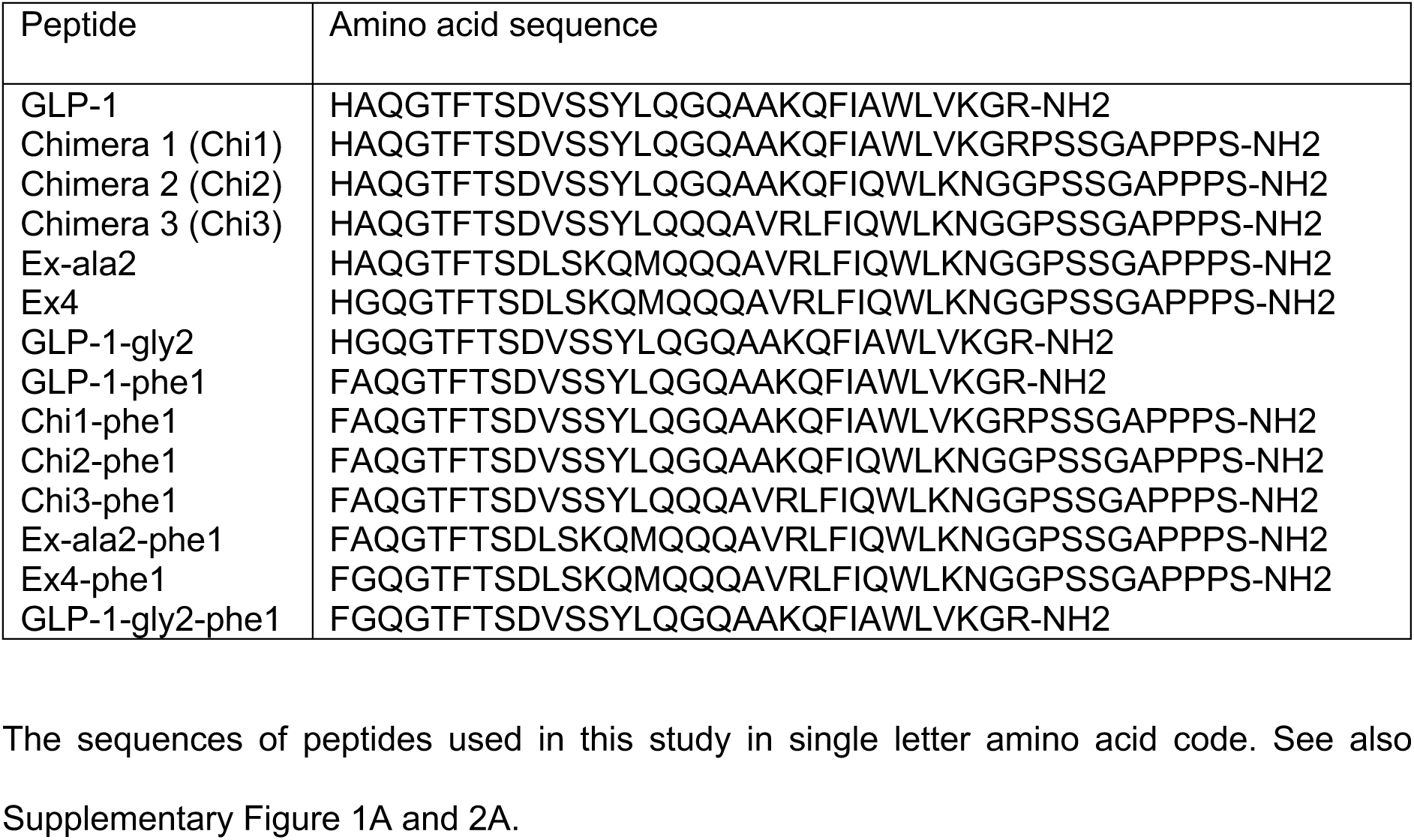
Peptides used in this study.

Equilibrium binding studies were performed in HEK293 cells expressing N-terminally SNAP-tagged GLP-1R (“HEK293-SNAP-GLP-1R”) labelled with the lanthanide time-resolved Förster resonance energy transfer (TR-FRET) donor SNAP-Lumi4-Tb, wherein the binding of unlabelled peptides was measured in competition with the fluorescent ligand exendin-4-FITC (14). Saturation binding of exendin-4-FITC was determined as part of each experiment (Supplementary Figure 1B). The most prominent finding was that the Chi3, Ex-ala2 and exendin-4 peptides showed increased binding affinity compared to GLP-1 itself, while GLP-1-gly2 displayed reduced affinity (Figure 1A, Table 2). Cyclic AMP (cAMP) and β-arrestin-2 recruitment responses were assessed in PathHunter CHO-K1-βarr2-EA-GLP-1R cells (Figure 1B, Table 2). Both Gly2 ligands (exendin-4 and GLP-1-gly2) displayed moderately but consistently reduced efficacy for β-arrestin-2 recruitment, in line with earlier work with exendin-4 (20). However, no ligand showed significant bias towards either pathway when analysed using the **τ**/K_A_ approach derived from the operational model of agonism (21) (Figure 1C).

**Figure 1.**
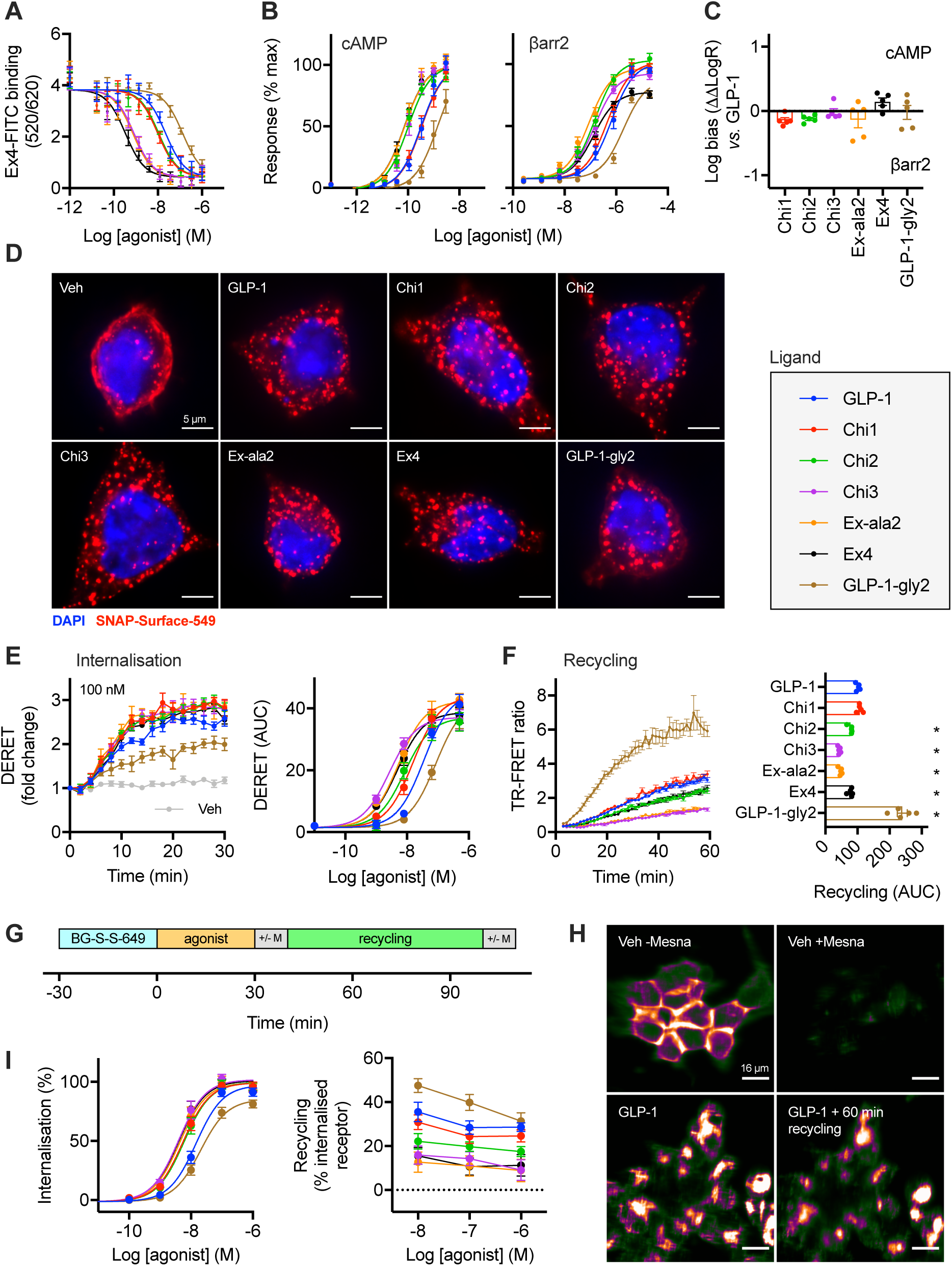
Binding, signalling and trafficking of chimeric GLP-1R agonist ligands. (**A**) Equilibrium binding studies in HEK293-SNAP-GLP-1R cells, showing TR-FRET-determined binding of 4 nM exendin-4-FITC in competition with indicated concentration of unlabelled agonist, *n*=5. See also Supplementary Figure 1B. (**B**) cAMP and β-arrestin-2 (βarr2) recruitment responses measured in parallel in CHO-K1-βarr2-EA-GLP-1R cells, *n*=5, with 3-parameter fits of pooled data shown. (**C**) Quantification of signal bias from data presented in (B) using ΔΔLog**τ**/K_A_ method, depicted relative to GLP-1, with statistical comparison performed by one-way randomised block ANOVA with Dunnett’s test using ΔLog**τ**/K_A_ values (i.e. prior to normalisation to GLP-1), with no ligand found to be significantly biased relative to GLP-1. (**D**) Deconvolved widefield microscopy maximum intensity projection images of HEK293-SNAP-GLP-1R cells labelled with SNAP-Surface-549 prior to stimulation with 1 µM agonist for 30 minutes, representative images of *n*=3 independent experiments. Scale bar = 5 µm. (**E**) Real time SNAP-GLP-1R internalisation in HEK293-SNAP-GLP-1R cells, measured by DERET, with response to 100 nM agonist shown as well as concentration responses representing AUC from traces shown in Supplementary Figure 1C, *n*=5, with 3-parameter fits of pooled data. (**F**) SNAP-GLP-1R recycling measured by TR-FRET in CHO-K1-SNAP-GLP-1R cells after BG-S-S-Lumi4-Tb labelling, stimulation with 100 nM agonist for 30 minutes, cleavage of residual surface BG-S-S-Lumi4-Tb, and 60 minutes recycling in the presence of LUXendin645 (10 nM), *n*=5, with AUC compared by one-way randomised block ANOVA with Dunnett’s test *versus* GLP-1. (**G**) Principle of high content microscopy assay to measure GLP-1R internalisation and recycling. (**H**) Example images taken from high content microscopy assay, showing the effect of Mesna on vehicle-treated cells (to demonstrate the efficiency of surface cleavage of BG-S-S-649) and of cells treated with 1 µM GLP-1 and then sequential Mesna application before and after recycling period (demonstrating signal intensity reduction from the same cell population reflecting cleavage of recycled surface receptor), scale bar = 16 µm. (**I**) Internalisation and recycling responses of chimeric GLP-1R ligands measured in HEK293-SNAP-GLP-1R cells by high content microscopy, 3-parameter fits of pooled internalisation data shown, *n*=11 for internalisation and *n*=5 for recycling. Data represented as mean ± SEM, with individual replicates shown in some cases.

**Table 2.** Binding and signalling parameter estimates for chimeric peptides.

| | Equilibrium binding | cAMP | | | β-arrestin-2 | | |
| --- | --- | --- | --- | --- | --- | --- | --- |
| | pK<sub>i</sub> (M) | pEC<sub>50</sub> (M) | E<sub>max</sub> (%) | Log $\tau$/K<sub>A</sub> (M) | pEC<sub>50</sub> (M) | E<sub>max</sub> (%) | Log $\tau$/K<sub>A</sub> (M) |
| GLP-1 | 7.8 ± 0.1 | 9.5 ± 0.1 | 100 | 9.5 ± 0.1 | 6.2 ± 0.1 | 102 ± 5 | 6.2 ± 0.1 |
| Chi1 | 8.2 ± 0.2 | 9.5 ± 0.1 | 100 | 9.5 ± 0.1 | 6.4 ± 0.1 | 102 ± 6 | 6.4 ± 0.1 |
| Chi2 | 8.2 ± 0.1 * | 10.0 ± 0.1 * | 100 | 10.0 ± 0.1 * | 6.9 ± 0.1 * | 104 ± 6 | 6.9 ± 0.1 * |
| Chi3 | 9.4 ± 0.1 * | 10.0 ± 0.1 * | 100 | 10.0 ± 0.1 * | 6.9 ± 0.2 * | 93 ± 5 * | 6.8 ± 0.1 * |
| Ex4-ala2 | 9.4 ± 0.1 * | 10.1 ± 0.1 * | 100 | 10.1 ± 0.1 * | 7.1 ± 0.2 * | 97 ± 5 | 6.8 ± 0.1 * |
| Ex4 | 9.7 ± 0.1 * | 10.2 ± 0.1 * | 100 | 10.1 ± 0.1 * | 6.9 ± 0.1 * | 77 ± 3 * | 7.0 ± 0.1 * |
| GLP-1-gly2 | 7.1 ± 0.1 * | 8.8 ± 0.1 * | 100 | 8.9 ± 0.1 * | 5.7 ± 0.1 * | 87 ± 4 * | 5.7 ± 0.1 * |

Due to the important biological role of GLP-1R endocytosis (8,9,14), we imaged SNAP-GLP-1R endosomal uptake in HEK293 cells via surface SNAP-labelling prior to agonist treatment (Figure 1D). At a single high dose, treatment with all ligands resulted in extensive and similar GLP-1R internalisation. We used diffusion-enhanced resonance energy transfer (DERET) (22) to monitor SNAP-GLP-1R internalisation in real-time at a range of doses (Figure 1E, Supplementary Figure 1C), which confirmed that all ligands were capable of inducing a high level of GLP-1R endocytosis at a maximal dose, albeit with significant differences in potency (Table 3). As post-endocytic sorting is an important factor regulating the surface levels of GLP-1R at steady state agonist stimulation (14), we also measured GLP-1R recycling using a cleavable form of SNAP-Lumi4-Tb (“BG-S-S-Lumi4-Tb”). In this assay, the reducing agent Mesna is used to remove residual GLP-1R labelling at the cell surface after an initial agonist-mediated internalisation step, with re-emergence of labelled GLP-1Rs at the cell surface measured in real time after agonist washout through binding to the far red fluorescent GLP-1R antagonist LUXendin645 (23), as previously described using a different acceptor ligand (24). LUXendin645 showed a large and rapid signal increases on binding to Lumi4-Tb-labelled SNAP-GLP-1R (Supplementary Figure 1D), indicating its suitability as a TR-FRET acceptor. Using this approach, marked differences in recycling rate *versus* GLP-1 were observed for GLP-1-gly2 (faster), as well as for Chi2, Chi3, exendin-4 and Ex-ala2, all of which recycled more slowly than GLP-1 itself (Figure 1F). These findings were confirmed by microscopy (Supplementary Figure 1E).

**Table 3.**
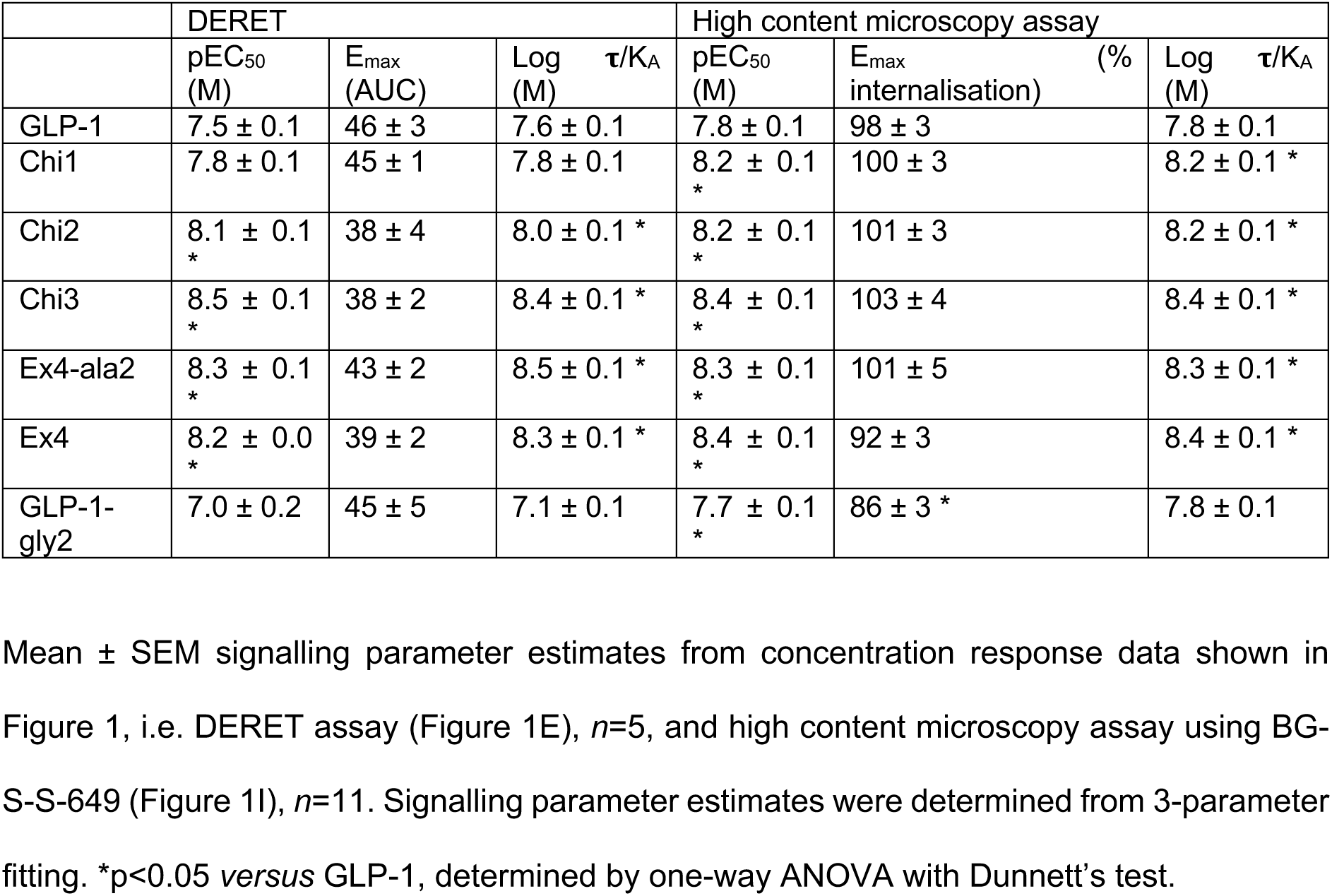
GLP-1R internalisation concentration response parameter estimates for chimeric ligands in HEK293-SNAP-GLP-1R cells.

|  | DERET |  |  | High content microscopy assay |  |  |
| --- | --- | --- | --- | --- | --- | --- |
| | pEC<sub>50</sub><br>(M) | E<sub>max</sub><br>(AUC) | Log $\tau$/K<sub>A</sub><br>(M) | pEC<sub>50</sub><br>(M) | E<sub>max</sub><br>internalisation (%) | Log $\tau$/K<sub>A</sub><br>(M) |
| GLP-1 | 7.5 ± 0.1 | 46 ± 3 | 7.6 ± 0.1 | 7.8 ± 0.1 | 98 ± 3 | 7.8 ± 0.1 |
| Chi1 | 7.8 ± 0.1 | 45 ± 1 | 7.8 ± 0.1 | 8.2 ± 0.1<br>* | 100 ± 3 | 8.2 ± 0.1 * |
| Chi2 | 8.1 ± 0.1<br>* | 38 ± 4 | 8.0 ± 0.1 * | 8.2 ± 0.1<br>* | 101 ± 3 | 8.2 ± 0.1 * |
| Chi3 | 8.5 ± 0.1<br>* | 38 ± 2 | 8.4 ± 0.1 * | 8.4 ± 0.1<br>* | 103 ± 4 | 8.4 ± 0.1 * |
| Ex4-ala2 | 8.3 ± 0.1<br>* | 43 ± 2 | 8.5 ± 0.1 * | 8.3 ± 0.1<br>* | 101 ± 5 | 8.3 ± 0.1 * |
| Ex4 | 8.2 ± 0.0<br>* | 39 ± 2 | 8.3 ± 0.1 * | 8.4 ± 0.1<br>* | 92 ± 3 | 8.4 ± 0.1 * |
| GLP-1-gly2 | 7.0 ± 0.2 | 45 ± 5 | 7.1 ± 0.1 | 7.7 ± 0.1<br>* | 86 ± 3 * | 7.8 ± 0.1 |

We also developed a high-content microscopy assay to sequentially measure GLP-1R internalisation and recycling at multiple doses using the Mesna-cleavable SNAP-tag probe BG-S-S-649 (Figure 1G). This assay represents a higher throughput adaption of an earlier flow cytometry assay (9). The far red DY-649 fluorophore was considered particularly suitable as it avoids the higher autofluorescence of plastic microplates at lower wavelengths. Sequential applications of Mesna after internalisation and recycling allowed determination of receptor distribution from the same fields of view within each well. Example images showing the effect of Mesna application before and after recycling are shown in Figure 1H (note that quantification was performed from several fields-of-view per well, representing many hundreds of cells). Results were broadly consistent with those obtained with the TR-FRET based assays (Figure 1I, Table 3).

Finally, coupling of occupancy to signalling and endocytosis was compared by subtracting log **τ**/K_A_ estimates from affinity measures and normalising to GLP-1 as the reference ligand (Supplementary Figure 1F). This analysis showed that the highest affinity ligands Chi3, Ex-ala2 and exendin-4 were less well coupled to functional responses than GLP-1, i.e. the greater affinity of these ligands did not result in commensurate increases in signalling.

These results highlight how sequence divergence between GLP-1 and exendin-4 can influence the binding affinity, signalling potency and trafficking characteristics of GLP-1R agonist ligands.

### 2.2 N-terminal substitution differentially affects binding, signalling and trafficking characteristics of chimeric peptides

Substitution of the exendin-4 N-terminal His to Phe results in reduced β-arrestin recruitment and internalisation (14). Nevertheless, structural differences in the host peptide might lead to changes to agonist orientation, N-terminal flexibility, and receptor interactions formed by agonist residues in the immediate vicinity, modulating the effects of the Phe1 substitution. We pharmacologically evaluated Phe1 analogues (Table 1, Supplementary Figure 2A) of the chimeric peptide series, first measuring binding affinities at equilibrium in HEK-SNAP-GLP-1R cells (Supplementary Figure 2B, Table 4). All Phe1 ligands showed reduced affinity compared to their His1 counterparts, included in the assay for parallel comparisons. Signalling parameters for each ligand were assessed in CHO-K1-βarr2-EA-GLP-1R cells (Figure 2A, Table 4). The most notable finding was a significantly reduced efficacy for β-arrestin-2 recruitment for all Phe1 ligands, particularly when Phe1 was introduced to both exendin-4 and GLP-1-gly2, with a more moderate effect for ligands containing Ala2 instead of Gly2. Signal bias of each ligand relative to GLP-1 is displayed in the heatmap, demonstrating that Phe1 analogues incorporating more of the exendin-4 sequence show progressively greater bias towards cAMP signalling, with Ex-phe1 being the most prominent example. Bias estimates are also displayed in an alternative format in Supplementary Figure 2C. Coupling of occupancy to signalling was determined from subtraction of log **τ**/K_A_ estimates for each pathway from pK_i_ for each ligand (Supplementary Figure 2D). This analysis suggests that introduction of Phe1 to Chi2, Chi3, Ex-ala2 and exendin-4 results in improved coupling to cAMP production, with lesser effects on β-arrestin-2 recruitment.

**Figure 2.**
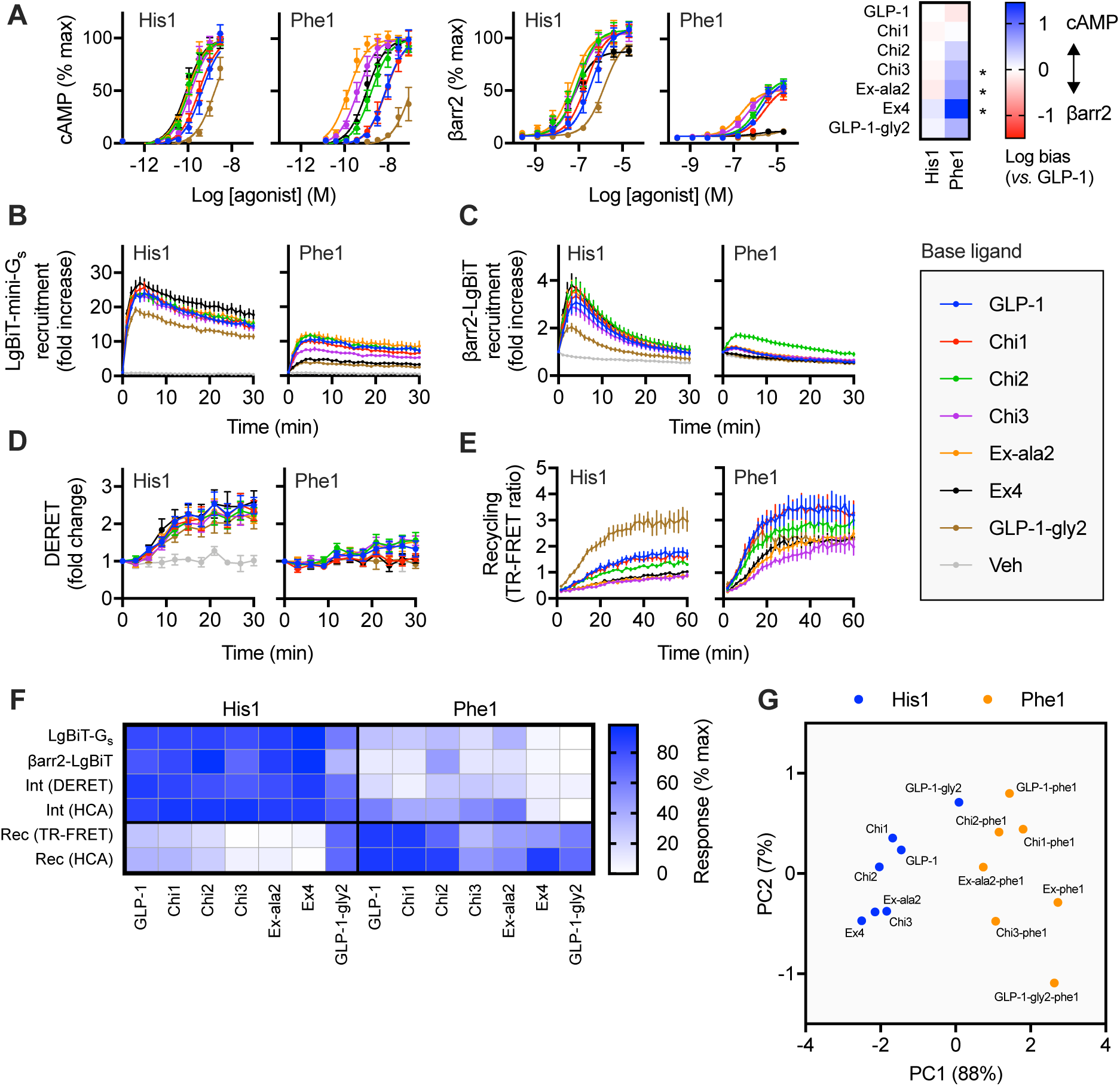
Pharmacological characterisation of N-terminally substituted chimeric GLP-1R agonists. (**A**) cAMP and β-arrestin-2 recruitment responses in CHO-K1-βarr2-EA-GLP-1R cells, *n*=5, with 3-parameter fits of pooled data shown; the heatmap shows signal bias quantified using the ΔΔLog**τ**/K_A_ method bias relative to GLP-1; statistical comparison was performed by one-way randomised block ANOVA with Sidak’s test to compare bias for each His1 / Phe1 ligand pair using ΔLog**τ**/K_A_ values (i.e. prior to normalisation to GLP-1). (**B**) NanoBiT measurement of LgBiT-mini-G_s_ recruitment to GLP-1R-smBiT in HEK293T cells after stimulation with a saturating (1 µM) concentration of ligand, *n*=5. (**C**) As for (B), but for βarr2-LgBit. (**D**) As for (B), but for GLP-1R internalisation in HEK293-SNAP-GLP-1R cells measured by DERET. (**E**) SNAP-GLP-1R recycling measured by TR-FRET in CHO-K1-SNAP-GLP-1R cells after BG-S-S-Lumi4-Tb labelling, stimulation with 1 µM agonist for 30 minutes, cleavage of residual surface BG-S-S-Lumi4-Tb, and 60 minutes recycling in the presence of LUXendin645 (10 nM), *n*=5. (**F**) Representation of data shown in (B), (C), (D), (E), Supplementary Figures 2I and 2J (internalisation [“Int”] and recycling [Rec] measured by high content microscopy analysis [HCA]), showing the mean response of each ligand with normalisation to the minimum and maximum ligand response in each assay repeat. (**G**) Principal component analysis of His1 and Phe1 ligands, derived from single dose response data from (B), (C), and Supplementary Figures 2I and 2J. *p<0.05 by statistical test indicated in the text. Data represented as mean ± SEM.

**Table 4.** Binding and signalling parameter estimates for His1- and Phe1-containing chimeric ligands.

| | Equilibrium binding | cAMP | | | β-arrestin-2 | | |
| --- | --- | --- | --- | --- | --- | --- | --- |
| | pK<sub>i</sub> (M) | pEC<sub>50</sub> (M) | E<sub>max</sub> (%) | Log $\tau$/K<sub>A</sub> (M) | pEC<sub>50</sub> (M) | E<sub>max</sub> (%) | Log $\tau$/K<sub>A</sub> (M) |
| GLP-1 | 8.5 ± 0.1 | 9.4 ± 0.1 | 100 | 9.3 ± 0.1 | 6.4 ± 0.2 | 111 ± 7 | 6.4 ± 0.1 |
| GLP-1-phe1 | 6.9 ± 0.1 *, # | 8.1 ± 0.2<br>*, # | 100 | 8.1 ± 0.1<br>*, # | 5.7 ± 0.1<br>*, # | 61 ± 7<br>*, # | 5.4 ± 0.1<br>*, # |
| Chi1 | 8.6 ± 0.1 | 9.5 ± 0.1 | 100 | 9.5 ± 0.1 | 6.6 ± 0.1 | 108 ± 6 | 6.6 ± 0.1 |
| Chi1-phe1 | 6.9 ± 0.1 *, # | 8.1 ± 0.2<br>*, # | 100 | 8.1 ± 0.2<br>*, # | 5.6 ± 0.0<br>*, # | 55 ± 7<br>*, # | 5.2 ± 0.1<br>*, # |
| Chi2 | 8.7 ± 0.2 | 10.0 ± 0.1 * | 100 | 9.9 ± 0.1 * | 7.0 ± 0.1 * | 108 ± 6 | 7.0 ± 0.1 * |
| Chi2-phe1 | 6.4 ± 0.1 *, # | 8.8 ± 0.2<br>*, # | 100 | 8.7 ± 0.2<br>*, # | 5.9 ± 0.0<br>*, # | 62 ± 5<br>*, # | 5.5 ± 0.1<br>*, # |
| Chi3 | 9.5 ± 0.1 * | 9.9 ± 0.1 * | 100 | 9.8 ± 0.1 * | 7.0 ± 0.1 * | 105 ± 7 | 7.0 ± 0.1 |
| Chi3-phe1 | 8.3 ± 0.1 # | 9.4 ± 0.2 # | 100 | 9.3 ± 0.2 # | 6.3 ± 0.1 # | 53 ± 5<br>*, # | 5.9 ± 0.1 # |
| Ex-ala2 | 9.8 ± 0.1 * | 10.0 ± 0.1 * | 100 | 10.0 ± 0.1 * | 7.3 ± 0.1 * | 105 ± 8 | 7.2 ± 0.1 * |
| Ex-ala2-phe1 | 8.4 ± 0.1 # | 9.8 ± 0.1 * | 100 | 9.8 ± 0.1 * | 6.7 ± 0.1 # | 51 ± 5<br>*, # | 6.2 ± 0.0 # |
| Ex4 | 9.9 ± 0.1 * | 10.0 ± 0.1 * | 100 | 10.1 ± 0.1 * | 7.2 ± 0.1 * | 87 ± 4 | 7.0 ± 0.1 |
| Ex-phe1 | 7.8 ± 0.1 *, # | 9.0 ± 0.2<br>*, # | 100 | 8.9 ± 0.2<br>*, # | 6.0 ± 0.2 # | 12 ± 1<br>*, # | 4.6 ± 0.2<br>*, # |
| GLP-1-gly2 | 7.2 ± 0.1 * | 8.8 ± 0.1 * | 100 | 8.8 ± 0.1 * | 5.8 ± 0.0 * | 101 ± 5 | 5.8 ± 0.0 * |
| GLP-1-gly2-phe1 | 6.5 ± 0.2 *, # | 7.0 ± 0.2<br>*, # | 100 | 7.1 ± 0.2<br>*, # | 4.8 ± 0.2<br>*, # | 16 ± 1<br>*, # | 3.6 ± 0.3<br>*, # |

To gain further insights into signalling differences between His1 and Phe1 ligands, we used NanoBiT complementation (25) to measure recruitment of both catalytically inactive mini-G_s_ protein (26) and β-arrestin-2 to GLP-1R. Here, complementary elements of the 19.1 kDa nanoluciferase enzyme have been appended to the GLP-1R C-terminus and the mini-G_s_ N-terminus or β-arrestin-2 C-terminus, allowing monitoring of dynamic changes in G_s_ protein and β-arrestin-2 recruitment. A high ligand concentration (1 µM) was specifically chosen to ensure a high degree of receptor occupancy [at least 70% for the lowest affinity agonist Chi2-Phe1 according to the law of mass action (27)] in order to provide efficacy data without requiring full concentration responses. Here, all Phe1 ligands demonstrated reduced efficacy compared to His1 equivalents for both mini-G_s_ (Figure 2B, Supplementary Figure 2E) and β-arrestin-2 recruitment (Figure 2C, Supplementary Figure 2F). GLP-1R internalisation was also measured by DERET, with all Phe1 ligands displaying severely reduced endocytic tendency (Figure 2D, Supplementary Figure 2G). Phe1 ligands also recycled faster than their His1 equivalents in all cases, as detected using TR-FRET (Figure 2E, Supplementary Figure 2H), except for GLP-1-gly2-phe1, for which the His1 version also shows rapid recycling. High content microscopy showed a similar pattern in both internalisation and recycling (Supplementary Figure 2I, J).

To allow comparison of ligand characteristics, responses for each assay are compared in the heatmap shown in Figure 2F after normalisation to the most efficacious ligand on a per-assay basis. This highlights how the Phe1 group of ligands show lower efficacy for each of the signalling and internalisation readouts. The most dramatic impact is observed within (Gly2-containing) exendin-4 and GLP-1-gly2, in keeping with a similar finding for β-arrestin-2 efficacy using the PathHunter system (Figure 2A), and confirmed by expressing the response of each Phe1 agonist as a percentage of that of its His1 counterpart (Supplementary Figure 2K). Recycling typically displayed the inverse pattern, although more overlap was observed between the Phe1 and His1 groups, e.g. Chi3-phe1 and Ex-ala2-phe1-induced recycling rates approached those of the slower His1 ligands. Further analysis revealed a relationship between GLP-1R binding affinity and recycling rate, which was most apparent for agonists with moderate affinity (Supplementary Figure 2L). Principal component analysis (28) was used to visually represent clustering of agonists showing similar signalling and trafficking characteristics (Figure 2G). Phe1 were clearly distinguished from His1 ligands within the first principal component (PC1, accounting for 88% of the total variance). Of note, the exendin-4 / Ex-phe1 pair were separated to the greatest degree within PC1.

These results highlight how the Phe1 substitution can markedly affect the pharmacology of GLP-1R ligands, but to varying degrees dependent on other ligand sequence features.

### 2.3 Effect of chimeric GLP-1RA peptides in beta cells

Cellular context can influence the manifestations of signal bias (29). A major site of action for native GLP-1 and its therapeutic mimetics is the pancreatic beta cell, where it is coupled to potentiation of insulin secretion (6). We therefore performed further assessments of His1 and Phe1 peptides in INS-1 832/3 cells (30), a clonal beta cell line of rat origin. cAMP signalling responses were consistent with the patterns observed in CHO-K1-βarr2-EA-GLP-1R cells, with reduced potency observed with all Phe1 *versus* equivalent His1 ligands (Figure 3A, Table 5). SNAP-GLP-1R endocytosis following treatment with each agonist was monitored in INS-1 832/3 cells by DERET (Figure 3B, Supplementary Figure 3A) and confirmed by microscopy (Figure 3C). All Phe1 ligands led to less internalisation, with Gly2-containing Ex-phe1 and GLP-1-gly2-phe1 showing virtually undetectable DERET responses. The DERET assay also showed approximately twice as fast GLP-1R uptake in INS-1 832/3 compared to HEK293-SNAP-GLP-1R cells (i.e. Figure 2D), and that the Phe1 ligand responses in the beta cell model (except for Ex-phe1 and GLP-1-gly2-phe1) were less diminished compared with the equivalent His1 ligand response (see Supplementary Table 1 for kinetic comparisons). High content microscopy also showed reduced internalisation with Phe1 ligands, especially for Ex-phe1 and GLP-1-gly2-phe1 (Supplementary Figure 3B), and further highlighted how Phe1 agonist-mediated GLP-1R internalisation was less reduced in the beta cell model (Supplementary Figure 3C). Phe1 ligands recycled faster than His1 equivalents (Supplementary Figure 3D), with higher affinity Chi3-phe1 and Ex-ala2-phe1 showing the slowest recycling rates among the Phe1 ligands.

**Figure 3.**
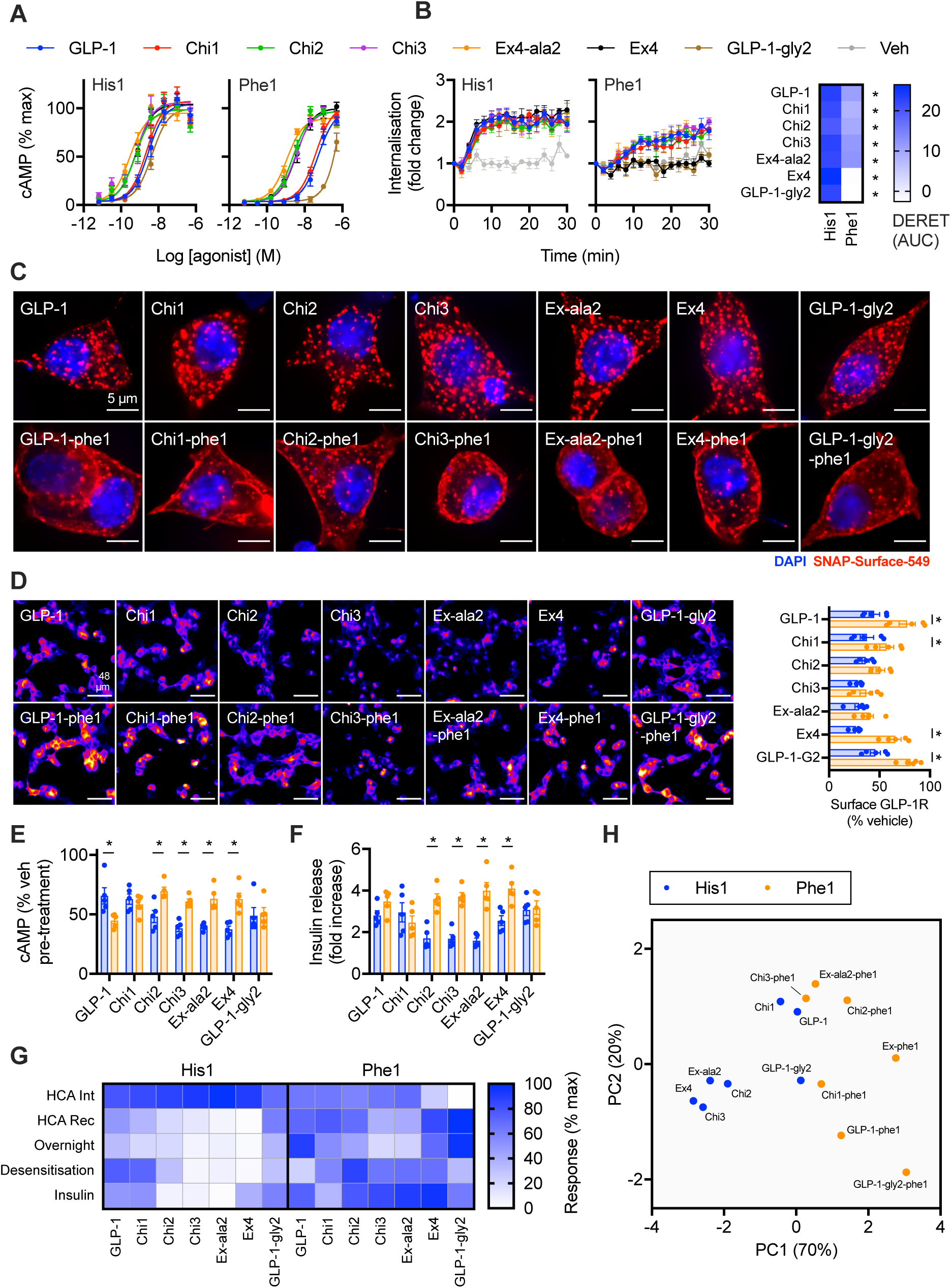
Effects of biased chimeric peptides in beta cells. (**A**) cAMP responses in INS-1 832/3 cells with endogenous GLP-1R expression stimulated for 10 minutes in the presence of 500 µM IBMX, *n*=5. (**B**) GLP-1R internalisation measured by DERET in INS-1 832/3 GLP-1R^-/-^ cells stimulated with 1 µM agonist, *n*=5, with quantification of AUC shown on the heatmap and statistically compared by one-way randomised block ANOVA with Sidak’s test for each His1 *versus* Phe1 ligand pair. (**C**) Widefield microscopy images of SNAP-GLP-1R cells labelled with SNAP-Surface-549 prior to stimulation with 1 µM agonist for 30 minutes, representative images of *n*=3 independent experiments. Scale bar = 5 µm. (**D**) Representative images showing GLP-1R surface expression levels detected by SNAP-Surface-649 labelling performed after 16-hour exposure to 1 µM agonist, scale bar = 48 µm, with quantification of *n*=5 experiments and comparison by one-way randomised block ANOVA with Sidak’s test for each His1 *versus* Phe1 ligand pair. (**E**) Homologous desensitisation experiment in wild-type INS-1 832/3 treated for 16 hours with 1 µM agonist, followed by washout, 1-hour recovery, and re-challenge with 100 nM GLP-1 + 500 µM IBMX, normalised to response to vehicle pre-treated cells, *n*=5, one-way randomised block ANOVA with Sidak’s test for each His1 *versus* Phe1 ligand pair. (**F**) Insulin secretion from wild-type INS-1 832/3 cells treated with 11 mM glucose ± 1 µM agonist for 16 hours, expressed relative to vehicle, *n*=5, one-way randomised block ANOVA with Sidak’s test for each His1 *versus* Phe1 ligand pair. (**G**) Representation of data shown in Supplementary Figures 3B and 3D, and Figures 3D-F, showing the mean response of each ligand with normalisation to the minimum and maximum ligand response in each assay repeat. (**H**) Princip component analysis of His1 and Phe1 ligands, derived from single dose response data from as represented in (G). *p<0.05 by statistical test indicated in the text. Data represented as mean ± SEM, with individual replicates shown in some cases.

**Table 5.**
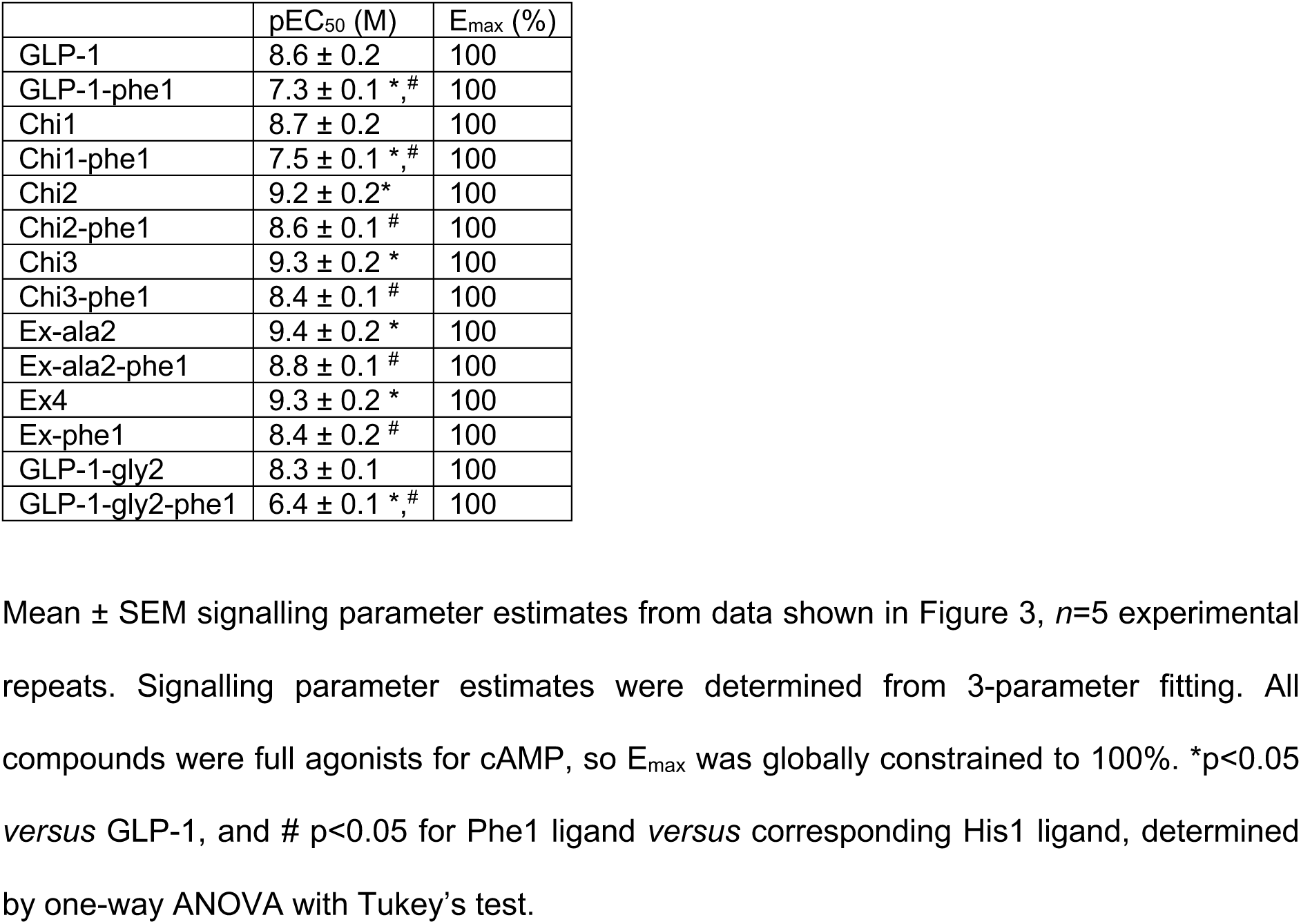
cAMP signalling parameter estimates of His1- and Phe1-containing chimeric ligands in INS-1 832/3 cells.

|  | pEC<sub>50</sub> (M) | E<sub>max</sub> (%) |
| --- | --- | --- |
| GLP-1 | 8.6 ± 0.2 | 100 |
| GLP-1-phe1 | 7.3 ± 0.1 *, # | 100 |
| Chi1 | 8.7 ± 0.2 | 100 |
| Chi1-phe1 | 7.5 ± 0.1 *, # | 100 |
| Chi2 | 9.2 ± 0.2* | 100 |
| Chi2-phe1 | 8.6 ± 0.1 # | 100 |
| Chi3 | 9.3 ± 0.2 * | 100 |
| Chi3-phe1 | 8.4 ± 0.1 # | 100 |
| Ex-ala2 | 9.4 ± 0.2 * | 100 |
| Ex-ala2-phe1 | 8.8 ± 0.1 # | 100 |
| Ex4 | 9.3 ± 0.2 * | 100 |
| Ex-phe1 | 8.4 ± 0.2 # | 100 |
| GLP-1-gly2 | 8.3 ± 0.1 | 100 |
| GLP-1-gly2-phe1 | 6.4 ± 0.1 *, # | 100 |

As the functional effects of biased GLP-1R ligands such as Ex-phe1 tend to emerge after prolonged stimulations (14), we tested the effect of a 16-hour exposure to each ligand on beta cell GLP-1R distribution by surface SNAP-labelling at the end of the exposure period. Note that this approach detects both recycled and newly synthesised GLP-1Rs, unlike the recycling assays used earlier in the present study in which only receptor labelled prior to stimulation is measured. Nevertheless, results were generally consistent with the trafficking characteristics of each ligand, with Phe1 agonists preserving more surface GLP-1R at the end of the exposure period than His1 agonists, with the exception of the Chi3 / Chi3-phe1 and Ex-ala2 / Ex-ala2-phe1 pairs (Figure 3D). To test the functional implications of these trafficking differences, we assessed for homologous GLP-1R desensitisation with each ligand by pre-treating for 16 hours, washing, and allowing a 1-hour recycling period before re-challenging with a fixed dose of GLP-1. Here, prior treatment with the slow recycling, high affinity His1 ligands Chi2, Chi3, Ex-ala2 and exendin-4 led to greater desensitisation, whereas the equivalent Phe1 analogues allowed cells to retain more responsiveness (Figure 3E). Similarly, continuous exposure to the same set of Phe1 ligands led to higher levels of cumulative insulin secretion (Figure 3F), suggesting that sustained insulinotropism is partly controlled by GLP-1R trafficking. To check that these findings were not artefacts of differences in potency, we also measured sustained insulin secretion with GLP-1, GLP-1-phe1, exendin-4 and Ex-phe1 over a range of doses, comparing these findings with previously determined acute cAMP potency measurements (Supplementary Figure 3E, F). This indicated that both GLP-1-like ligands displayed a relative reduction in potency for sustained insulin secretion *versus* acute cAMP potency compared to the exendin peptides, likely to indicate accelerated ligand degradation, but also that the high ligand dose used in earlier experiments remained maximal during prolonged exposure (and is therefore indicative of efficacy).

Thus, Phe1 ligands were generally characterised by slower internalisation, faster recycling, reduced net loss of surface GLP-1R and desensitisation, as well as greater cumulative insulin secretion. However, comparison of the relative responses of each ligand in each assay indicated that trafficking characteristics did not entirely predict functional responses (Figure 3G). For example, within the Phe1 group of ligands, Chi3-phe1 and Ex-ala2-phe1 showed the greatest loss of surface receptor after overnight exposure, in keeping with their somewhat greater acute internalisation and reduced recycling; however, this was not associated with commensurate increases in desensitisation or reduced insulin secretion. Nevertheless, principal component analysis incorporating functional cAMP and insulin readouts as well as trafficking data again showed clear discrimination of Phe1 *versus* His1 peptides, with Ex-phe1 and exendin-4 showing the most marked difference within the first principal component (Figure 3H). A correlation matrix summarising the relationship between agonist parameters measured across the different cell lines used in this study is shown in Supplementary Figure 3E.

Thus, the overall pattern of biased GLP-1R action identified in HEK293 and CHO-K1 cells was at least partially replicated in a beta cell model. The functional impact of altered GLP-1R trafficking partly, but not entirely, explained the differences in sustained insulin secretion observed with the Phe1 ligands.

## 3 Discussion

In this study we evaluated a series of GLP-1R ligands with variable sequence homology to GLP-1 and exendin-4, focusing in particular on their effects on signal bias and GLP-1R trafficking. We previously found that the functional and therapeutic effects of exendin-4-derived peptides can be profoundly affected by N-terminal sequence modifications that preferentially alter β-arrestin recruitment, GLP-1R endocytosis and recycling (14). The present work builds on our earlier findings by assessing further ligand factors that influence these processes.

We developed a high content microscopy-based assay to measure GLP-1R internalisation and recycling in the same assay using a cleavable SNAP-tag-labelling far red fluorescent probe. This approach was adapted from earlier work by us (9) and others (31) in which similar fluorescent probes were used in lower throughput assays based on flow cytometry and confocal microscopy, respectively. Simultaneous handling of multiple ligands or ligand doses in multi-well plates, along with data acquisition from several fields of view at relatively lower magnification, enhances experimental reproducibility and statistical robustness compared to lower throughput methods (32). The cleavable BG-S-S-Lumi4-Tb TR-FRET method for measuring GLP-1R recycling provided generally concordant results with those obtained using the high content assay. One example where results did not entirely agree was for exendin-4 *versus* Ex-phe1 (Figure 2F), but it is not clear if this represents experimental variability, cell-type differences (HEK293 *versus* CHO-K1), or methodological variances, e.g. due to the more indirect method of detection with the TR-FRET assay. The latter method gives additional kinetic information not available from our implementation of the microscopy-based Mesna cleavage method, although fast-recycling events could in principle be studied by monitoring fluorescence loss in real time during continuous Mesna exposure (31). This was not attempted here as, for many of the slow-recycling ligands, the required exposure time to a reducing environment and non-physiological pH would lead to non-specific effects on cell behaviour.

We found clear binding, signalling and internalisation differences between GLP-1 and exendin-4, with affinity for the latter being approximately 5 times greater and potency (for most readouts) approximately 10 times greater. As the GLP-1 amino acid sequence was progressively replaced by that of exendin-4, starting at the C-terminus, both affinity and potency increased. Of note, the impact of adding the exendin-4 C-terminal extension, or “Trp-cage”, was small, consistent with other studies in which its truncation had little effect on exendin-4 signalling (33). It should be noted that, once occupancy differences are taken into account, exendin-4-like ligands show a comparative signalling deficit, as greater affinity did not fully translate into commensurate increases in signalling potency. A recent report describes the detection of two separate binding sites for exendin-4 on the GLP-1R extracellular domain (ECD), potentially in keeping with previous photoaffinity cross-linking data (34) and apparently responsible for impaired receptor oligomerisation compared to non-exendin-4 ligands (35). As GLP-1R dimerisation is required for full signalling efficacy (36), it is possible that this phenomenon partly explains our results. However, the full-length GLP-1R may not behave identically to the isolated ECD (37, 38). The Ala → Gly switch at position 2, seen in exendin-4 and GLP-1-gly2, was noted in the present study to modestly reduce signalling efficacy for β-arrestin-2 in both cases. This is in keeping with previous work (20) and potentially relevant to the signalling characteristics of Phe1 ligands (see below). Contrasting with this consistent signalling efficacy effect, affinity of GLP-1-gly2 was almost 10-fold reduced *versus* GLP-1, whereas exendin-4 showed similar binding characteristics to Ex-ala2. This highlights how binding affinity of GLP-1-like peptides depends more on interactions with the receptor core made by their N-terminal regions than for exendin-4-like ligands, for which the more helical mid-peptide regions and C-terminus allow stable interactions with the ECD (19).

None of the His1-containing ligands showed significant signal bias relative to GLP-1, although a trend for cAMP-preference was observed for exendin-4, in keeping with some (20) but not all (10) previous literature. Moreover, endocytosis potency differences generally followed the pattern established for cAMP and β-arrestin-2 signalling. GLP-1R recycling measured by two different methods highlighted clear differences in GLP-1-like *versus* exendin-4-like peptides across a wide range of doses, with the latter showing more gradual recycling. A similar pattern was found for GLP-1 *versus* exendin-4 in a previous study (39). In the latter work, the difference was suggested to be partly related to GLP-1 degradation by dipeptidyl dipeptidase-4 (DPP-4), which cleaves at the penultimate Ala residue close to the N-terminus and to which exendin-4 is resistant by virtue of the Ala → Gly switch at position 2. However, our observation of slow recycling with DPP-4-sensitive Ex-ala2 suggests that DPP-4 is unlikely to be a critical determinant of GLP-1R recycling. Interestingly, endothelin converting enzyme-1 (ECE-1) has recently been shown to play a role in the control of GLP-1R recycling and desensitisation (40), presumably due to intra-endosomal ligand degradation and subsequent re-routing of unbound GLP-1R to a recycling pathway. Due to its enhanced resistance to other endopeptidases such as neprilysin (41), exendin-4 is likely to be ECE-1-resistant, although this has not been confirmed experimentally. In conjunction with its higher binding affinity, this could potentially contribute to more persistent intra-endosomal GLP-1R occupancy, resulting in accentuated targeting to post-endocytic degradative pathways. This possibility could be amenable to future exploration.

Due to the beneficial effects of exendin-phe1 (14), we wished to determine whether sequence changes elsewhere in the peptide would modulate the effects of the Phe1 N-terminal substitution. Concentration response experiments in CHO-K1 cells revealed reduced efficacy for β-arrestin-2 with Phe1 ligands in all cases, although the effect was most apparent with the Gly2-containing Ex-phe1 and GLP-1-gly2-phe1, for which β-arrestin-2 recruitment was virtually undetectable, even in the inherently amplified DiscoverX system. The efficacy reduction for the combination of Phe1 and Gly2 appeared to be at least additive. Interestingly, pathway bias analysis did not universally show that this reduction in β-arrestin-2 efficacy translated to significant bias in favour of cAMP, with only Chi3-phe1, Ex-ala2-phe1 and Ex-phe1 showing statistically significant changes compared to their His1 counterparts. This may partly relate to the inherently increased imprecision for low efficacy agonists (42); alternatively, it could imply that relatively high affinity (as is the case for these three Phe1 ligands) is required for a significant degree of bias. The NanoBiT complementation approach provides a means to compare dynamic G protein *versus* β-arrestin-2 recruitment events without the caveats of adenylate cyclase amplification of cAMP responses or irreversible enzyme complementation in the DiscoverX system. Here, Phe1 ligands were found in all cases to show lower efficacy for both pathways. This reduction, however, appeared more marked for β-arrestin-2 recruitment, suggesting efficacy-driven bias in favour of G_s_. Again, the Gly2-containing Phe1 ligands showed the greatest signalling deficit of all, with small magnitude responses even for G_s_ recruitment (which are, nevertheless, sufficient to produce full cAMP responses in the context of adequate amplification by adenylate cyclase). It should be noted that mini-G protein recruitment responses do not fully recapitulate G protein activation dynamics, which have recently been studied using NanoBRET-based sensors (43) and would be an interesting future investigation for these ligands. The β-arrestin-2 recruitment response of Chi2-phe1 was greater than that of other Phe1 agonists, which was not observed in the DiscoverX assay; this observation remains unexplained.

GLP-1R recycling data from a wide range of His1- and Phe1-containing ligands provided further insights into the relationship between GLP-1RA binding affinity and post-endocytic targeting, as highlighted in our earlier study (14). In the present work, affinity predicted recycling rates for ligands within a pK_i_ range of approximately 1 – 100 nM. The fact that some ligands with presumably enhanced proteolytic stability but lower affinity (e.g. Ex-ala2-phe1) were found to recycle faster than those with reduced stability but higher affinity (e.g. GLP-1) argues against intra-endosomal peptide degradation being the dominant factor influencing GLP-1R recycling, but this speculation requires experimental verification.

We performed specific studies in the pancreatic beta cell-like INS-1 832/3 model to identify whether the pharmacological properties of the ligands tested here are potentially relevant to biologically important insulin release. Measures from HEK293 cells were generally replicated in the beta cell model, although some differences in the trafficking characteristics were apparent. In particular, endocytosis was faster and more extensive in INS-1 cells, with His1 ligands reaching peak DERET signal up to twice as fast as in the HEK293 model, and with a less marked suppression of internalisation with many Phe1 ligands also observed. Whilst the specific trafficking characteristics of GLP-1R in HEK293 cells are not in themselves physiologically relevant, it raises the possibility of cell-specific differences in GLP-1R trafficking in physiological systems, i.e. a form of “tissue bias” (29). It could be speculated that different GLP-1RAs might therefore behave differently in beta cells and anorectic neurons, for example, although further experiments using primary neurons and beta cells would be required to substantiate this. Moreover, there is current interest in using GPCR-targeting peptides to deliver cargo to metabolically important tissues (44–46) and, as such, tissue-specific GLP-1R endocytic characteristics might be relevant to the targeting efficiency for different cell types and organ systems.

As therapeutic GLP-1RAs have now been engineered for prolonged pharmacokinetics allowing weekly administration, we focused the final part of this study on the beta cell effects of sustained agonist exposure. We observed that the acute trafficking responses of each peptide were reasonably predictive of surface GLP-1R levels after prolonged treatment, with fast recycling generally associated with greater detection of surface GLP-1R at the end of the incubation period. The slow internalising / fast recycling compounds (such as Ex-phe1 and GLP-1-gly2-phe1) had some of the highest levels of GLP-1R remaining at the surface in this assay. Interestingly, in spite of almost complete SNAP-GLP-1R internalisation within 30 minutes of high efficacy His1 agonist stimulation detected using cleavable BG-S-S-probes earlier in this study, the post-stimulation labelling approach suggested GLP-1R surface levels remained at least 30% of those of in vehicle-treated cells despite prolonged stimulation period at a high agonist dose. This may partly represent GLP-1R recycling during the labelling period but could also represent delivery of newly synthesised or constitutively recycled GLP-1Rs to the cell surface, which would not be detected using the pre-stimulation labelling approach.

Interestingly, functional readouts in wild-type INS-1 832/3 cells, including homologous desensitisation and insulin secretion, did not perfectly recapitulate the pattern of GLP-1R observed with prolonged agonist exposure. One example is the comparison between His1 and Phe1 versions of Ex-ala2 and exendin-4; here, both Phe1 ligands showed clearly reduced homologous desensitisation and improved insulin secretion, yet surface GLP-1R levels with Ex-ala2-phe1 were no different to Ex-ala2-his1. Further complexities in the spatiotemporal control of signalling might contribute to this dichotomy. Additionally, differences between the cell systems used for trafficking (INS-1 832/3 GLP-1R^-/-^ cells with SNAP-GLP-1R overexpression) and functional studies (wild-type INS-1 832/3 cells with endogenous GLP-1R expression) could be relevant. Insertion of biorthogonal tags, e.g. SNAP or Halo, into the endogenous receptor genomic sequence (47) may allow these events to be studied without overexpression artefacts and should be used in the future. Fluorescent ligands can also be used to study endogenous GLP-1R dynamics (23), provided fluorophore bioconjugation to a relatively large number of peptides (as in this study) is feasible and does not interfere with ligand pharmacological properties.

In summary, this work provides a systematic evaluation of a panel of GLP-1RAs with varying homology to GLP-1 and exendin-4. We identified marked differences in signalling, endocytosis and trafficking characteristics, which may be informative for the development of improved GLP-1RAs for T2D and obesity. As well as the possible future studies suggested above, these agonists may be useful in combination with the wide range of mutant GLP-1R constructs that have been published (5,48,49) to gain a better understanding of specific ligand-receptor interactions important for specific GLP-1R behaviours.

## 4 Methods

### 4.1 Peptides and Reagents

Peptides were obtained from Wuxi Apptec and were at least 90% pure. Homogenous time-resolved fluorescence (HTRF) reagents, including cAMP Dynamic 2 kit, wide-range insulin HTRF kit, SNAP-Lumi4-Tb and cleavable BG-S-S-Lumi4-Tb, were obtained from Cisbio. PathHunter detection reagents were obtained from DiscoverX. SNAP-Surface-549 and -649 probes were obtained from New England Biolabs. Cleavable BG-S-S-649 probe was a gift from New England Biolabs. Cell culture reagents were obtained from Sigma and Thermo Fisher Scientific.

### 4.2 Cell Culture

HEK293T cells were maintained in DMEM supplemented with 10% FBS and 1% penicillin/streptomycin. HEK293-SNAP-GLP-1R cells, generated by stable transfection of pSNAP-GLP-1R (Cisbio), previously described in (50), were maintained in DMEM supplemented with 10% FBS, 1% penicillin/streptomycin and 1 mg/ml G418. PathHunter CHO-K1-βarr2-EA-GLP-1R cells were maintained in Ham’s F12 medium supplemented with 10% FBS, 1% penicillin/streptomycin, 1 mg/ml G418 and 0.4 mg/ml hygromycin B. CHO-K1-SNAP-GLP-1R cells (50) were maintained in DMEM supplemented with 10% FBS, 10 mM HEPES, 1% non-essential amino acids, 1% penicillin/streptomycin and 1 mg/ml G418. Wildtype INS-1 832/3 cells (a gift from Prof. Christopher Newgard, Duke University) (30), and INS-1 832/3 cells lacking endogenous GLP-1R after deletion by CRISPR/Cas9 (INS-1 832/3 GLP-1R^-/-^ cells, a gift from Dr Jacqueline Naylor, MedImmune, Astra Zeneca) (51), were maintained in RPMI-1640 with 11 mM glucose, 10 mM HEPES, 2 mM glutamine, 1 mM sodium pyruvate, 50 µM β-mercaptoethanol, 10% FBS, and 1% penicillin/streptomycin. INS-1 832/3 SNAP-GLP-1R cells were generated from INS-1 832/3 GLP-1R^-/-^ cells by stable transfection of pSNAP-GLP-1R and selection with G418 (1 mg/ml).

### 4.3 TR-FRET Equilibrium Binding Assays

Binding of test peptides was monitored in competition with the fluorescent agonist exendin-4-FITC, in which the FITC is conjugated to Lys12 of the native exendin-4 sequence (52). Assays were performed in HEK293-SNAP-GLP-1R cells labelled with SNAP-Lumi4-Tb (40 µM, 30 min, in complete medium) followed by washing and resuspension in HBSS with 0.1% BSA, and supplemented with metabolic inhibitors (20 mM 2-deoxyglucose and 10 mM NaN_3_) to maintain GLP-1R at the cell surface during the assay (53). Labelled cells were placed at 4°C before addition of a range of concentrations of test peptides prepared in HBSS with 0.1% BSA containing 4 nM exendin-4-FITC. A range of concentrations of exendin-4-FITC was also used to establish K_d_ by saturation binding analysis for the assay. Cells were then incubated for 24 hours at 4°C before reading by time-resolved Förster resonance energy transfer (TR-FRET) in a Flexstation 3 plate reader (Molecular Devices) using the following settings: λ_ex_ = 335 nm, λ_em_ = 520 and 620 nm, delay 50 μs, integration time 400 μs. Binding was quantified as the ratio of fluorescent signal at 520 nm to that at 620 nm, after subtraction of ratio obtained in the absence of FITC-ligands. IC_50_ values for test peptides were determined using the “one site – fit K_i_” model in Prism 8.0 (GraphPad Software), with the K_d_ result for exendin-4-FITC obtained using the “one site – specific binding” model for that experiment used to constrain the assay.

### 4.4 Acute cyclic AMP Assays

Cells were seeded into white 96-well half area plates and stimulated with the indicated concentration of agonist in serum-free medium. CHO-K1-βarr2-EA-GLP-1R and HEK293-SNAP-GLP-1R cells were treated for 30 minutes at 37°C without phosphodiesterase inhibitors. Wild-type INS-1 832/3 cells, after a 6-hour preincubation in low glucose (3 mM) complete medium, were stimulated for 10 minutes at 37°C with 500 µM 3-isobutyl-1-methylxanthine (IBMX). HTRF detection reagents (cAMP Dynamic 2 kit, Cisbio) were added at the end of the incubation, and the plate was read after a further 60-minute incubation by HTRF using a Spectramax i3x plate reader (Molecular Devices).

### 4.5 PathHunter β-arrestin Recruitment Assays

CHO-K1-βarr2-EA-GLP-1R cells were seeded into white 96-well half area plates and stimulated for 30 minutes at 37°C with the indicated concentration of agonist in serum-free Ham’s F12 medium. PathHunter detection reagents were added and luminescence readings taken after 60 minutes at room temperature using a Spectramax i3x plate reader.

### 4.6 Homologous Desensitisation Assay

INS-1 832/3 cells were seeded into 96 well tissue-culture treated plates previously coated with 0.1 % poly-D-lysine, in complete medium at 11 mM glucose. Ligands were promptly added without IBMX and cells were incubated for 16 hours overnight at 37°C. The next day, medium was removed, and cells washed three times in HBSS. After a 60-minute recycling period in complete medium, cells were washed once more, and complete medium containing 500 µM IBMX + GLP-1 (100 nM) was added for 10 minutes before cell lysis. Lysates were analysed for cAMP concentration by HTRF as above, and results expressed relative to the vehicle pre-treatment control cells.

### 4.7 Insulin secretion measurements

Wild-type INS-1 832/3 cells were exposed to a 6-hour preincubation in low glucose (3 mM) complete medium before the assay. Cells were seeded in suspension into complete medium with 11 mM glucose ± agonist and incubated for 16 hours at 37°C. Secreted insulin in the supernatant was analysed by HTRF (Insulin High Range kit, Cisbio) after dilution and normalised to the concentration in glucose-only treated wells.

### 4.8 NanoBiT Complementation Assays

The GLP-1R-SmBiT plasmid was generated following digestion of GLP-1R-Tango (Addgene plasmid # 66295, a gift from Bryan Roth), with AgeI and XbaI and ligation of a duplexed SmBiT sequence (SmBiT F: 5’-ccggtggtggatccggcggaggtgtgaccggctaccggctgttcgaggagattctgtaat-3’; SmBiT R: 5’-gatctaatgtcttagaggagcttgtcggccatcggccagtgtggaggcggcctaggtggt-3’). The assay was performed as previously described (24). HEK293T cells in 12-well plates were co-transfected with 0.05 µg each of GLP-1R-SmBiT and βarr2-LgBiT plasmid (Promega) plus 0.9 µg pcDNA3.1, or with 0.5 µg each of GLP-1R-SmBiT and LgBiT-mini-G_s_ plasmid (a gift from Prof Nevin Lambert, Medical College of Georgia) (26). The assay was performed 24 – 36 hours later. Cells were detached, resuspended in NanoGlo Live Cell Reagent (Promega) with furimazine (1:20 dilution) and seeded into white 96-well half area plates. Baseline luminescence was immediately recorded over 5 minutes at 37°C in a Flexstation 3 plate reader, and serially after agonist addition for 30 minutes.

### 4.9 Measurement of GLP-1R Internalisation by DERET

The assay was performed as previously described (50). Cells were labelled with SNAP-Lumi4-Tb (40 µM, 30 min, in complete medium), washed, and resuspended in HBSS with 24 µM fluorescein. TR-FRET signals at baseline and serially after agonist addition were recorded at 37°C using a Flexstation 3 plate reader using the following settings: λ_ex_ = 335 nm, λ_em_ = 520 and 620 nm, delay 400 μs, integration time 1500 μs. Receptor internalisation was quantified as the ratio of fluorescent signal at 620 nm to that at 520 nm, after subtraction of individual wavelength signals obtained from wells containing 24 µM fluorescein only.

### 4.10 TR-FRET GLP-1R Recycling Assay

The method was adapted from a previous description (24), with the main change being the use of LUXendin645 (23) as the TR-FRET acceptor to improve signal-to-noise ratio. Adherent CHO-K1-SNAP-GLP-1R cells in 96-well half area tissue culture-treated plates were labelled with BG-S-S-Lumi4-Tb (40 µM, 30 minutes in complete medium), followed by washing. Agonist treatments were applied in serum-free medium for 30 minutes at 37°C. The plate was then placed on ice, washed with cold HBSS followed by 5 minute application of cold Mesna (100 mM in alkaline TNE buffer) to cleave Lumi4-Tb from receptors remaining at the cell surface. After further washing in the cold, warm HBSS containing 10 nM LUXendin645 was then added and TR-FRET serially monitored at 616 nm and 665 nm. Reappearance at the plasma membrane of Lumi4-Tb-labelled SNAP-GLP-1R is thus determined as an increase in TR-FRET, expressed ratiometrically as signal intensity at 665 nm divided by that at 616 nm.

### 4.11 Imaging of Receptor Redistribution

Cells seeded overnight on coverslips were labelled for 30 minutes at 37°C with SNAP-Surface 549 (1 µM) to label surface SNAP-GLP-1Rs. After washing, treatments were applied for 30 minutes at 37°C. Where indicated, cells were washed in HBSS twice, incubated in complete medium for one hour to allow GLP-1R recycling, followed by a further wash in HBSS, and surface receptor was labelled with LUXendin645 (100 nM) for 5 minutes. Cells were fixed with 2% paraformaldehyde for 20 minutes at room temperature. Coverslips were mounted with Diamond Prolong antifade (Thermo Fisher) with DAPI and imaged using a modular microscope platform from Cairn Research incorporating a Nikon Ti2E, LED light source (CoolLED) and a 0.95 numerical aperture 40X air objective, or 1.45 numerical aperture 100X oil immersion objective. Where indicated, Z-stacks were acquired and deconvolved using Huygens software (SVI). Image galleries were generated using Fiji.

### 4.12 High Content Microscopy Internalisation / Recycling Assay

Cells were seeded into poly-D-lysine-coated, black 96-well plates to promote attachment and avoid cell loss during the wash steps. On the day of the assay, labelling was performed with BG-S-S-649 (1 µM), a surface-labelling SNAP-tag probe that, like BG-S-S-Lumi4-Tb, can be released on application of reducing agents such as Mesna (9, 31). After washing, treatments were applied for 30 minutes at 37°C in complete medium. Ligand was removed and cells washed with cold HBSS, and placed on ice for subsequent steps. Mesna (100 mM in alkaline TNE buffer, pH 8.6) or alkaline TNE without Mesna was applied for 5 minutes, and then washed with HBSS. Imaging of microplates was performed immediately without fixation, using the microscope system in Section 4.11 fitted with a 20X phase contrast objective, assisted by custom-written high content analysis software (54) implemented in Micro-Manager (55). A minimum of 6 images per well was acquired for both epifluorescence and transmitted phase contrast. On completion of imaging, HBSS was removed and replaced with fresh complete medium and receptor was allowed to recycle for 60 minutes at 37°C, followed by a second Mesna application to remove any receptor that had recycled to the membrane, and the plate was re-imaged as above. Internalised SNAP-GLP-1R was quantified at both time points as follows using Fiji: 1) phase contrast images were processed using PHANTAST (56) to segment cell-containing regions from background; 2) illumination correction of fluorescence images was performed using BaSiC (57); 3) fluorescence intensity was quantified for cell-containing regions. Agonist-mediated internalisation was determined as the mean signal for each condition normalised to signal from wells not treated with Mesna, after first subtracting non-specific fluorescence determined from wells treated with Mesna but no agonist. The percentage reduction in residual internalised receptor after the second Mesna treatment was considered to represent recycled receptor. Recycling was then expressed as a percentage relative to the amount of receptor originally internalised in the same well.

### 4.13 Post-Treatment Receptor Labelling Assay

INS-1 832/3 SNAP-GLP-1R cells were seeded into poly-D-lysine-coated, black 96-well plates and treatments were promptly applied. Wild-type INS-1 832/3 (without SNAP-tag) were used as a control to account for any contribution of non-specific labelling. After an 16 hour overnight incubation, cells were washed 3 times in HBSS and labelled for 1 hour at 37°C with 1 µM SNAP-Surface-649, before washing and imaging. Imaging settings and cell segmentation were performed as described in 4.12. Mean signal intensity from wild-type INS-1 832/3 cells was subtracted, allowing surface SNAP-GLP-1R expression after agonist treatment to be expressed relative to vehicle-treated INS-1 832/3 SNAP-GLP-1R cells.

### 4.14 Data Analysis and Statistics

Quantitative data were analysed using Prism 8.0. One biological replicate was treated as the average of technical replicates from an independently performed experiment. Intensity quantification from imaging data was performed on full depth 16-bit data after illumination correction; images are displayed with reduced dynamic range to highlight dim structures, with the same brightness and contrast settings consistently applied across matched images. Binding affinity calculations are described in Section 4.3. 3-parameter logistic fitting was performed for concentration responses analyses with constraints imposed as described in the table legends. For bias calculations, to reduce contribution of inter-assay variability and to avoid artefactual bias resulting from different activation kinetics of each pathway (58), cAMP and β-arrestin-2 assays were performed concurrently, with the same incubation time of 30 minutes; bias was determined by calculating transduction coefficients (21, 59); here, due to the matched design of our experiments, we calculated ΔΔlog(τ/K_A_) on a per-assay basis by normalising the log(τ/K_A_) for each ligand to the reference ligand (GLP-1) and then to the reference pathway (β-arrestin-2). Principal component analysis was performed using ClustVis (60) using the “SVD with imputation” method with unit variance scaling applied. ANOVA and t-test approaches were used for statistical comparisons. In experiments with a matched design, randomised block one-way ANOVA was used to compare treatments, with specific post-hoc tests indicated in the figure legends. Statistical significance was inferred when p<0.05. Data are represented as mean ± SEM throughout, with individual experimental replicates shown where possible.

---

**Supplementary Table 1.** SNAP-GLP-1R internalisation rates for each ligand in HEK293 and INS-1 832/3 cells.

| Ligand | T<sub>1/2</sub> (HEK293) | T<sub>1/2</sub> (INS-1 832/3) | Ratio |
| --- | --- | --- | --- |
| GLP-1 | 9.9 | 4.8 | 2.1 |
| GLP-1-phe1 | 16.9 | 10.3 | 1.6 |
| Chi1 | 10.4 | 5.6 | 1.9 |
| Chi1-phe1 | n.c. | 16.8 | n.c. |
| Chi2 | 10.0 | 5.6 | 1.8 |
| Chi2-phe1 | 17.2 | 9.0 | 1.9 |
| Chi3 | 10.2 | 5.0 | 2.0 |
| Chi3-phe1 | 17.4 | 13.4 | 1.3 |
| Ex-ala2 | 8.9 | 4.6 | 1.9 |
| Ex-ala2-phe1 | 16.4 | 10.1 | 1.6 |
| Ex4 | 8.9 | 4.6 | 1.9 |
| Ex4-phe1 | n.c. | n.c. | n.c. |
| GLP-1-gly2 | 10.9 | 5.0 | 2.2 |
| GLP-1-gly2-phe1 | n.c. | n.c. | n.c. |

Internalisation half-times were calculated by fitting a 4-parameter logistic curve to pooled DERET data expressed as fold-increase from baseline, with “basal response” constrained to a fixed value of 1. Ratio indicates the ratio of internalisation t<sub>1/2</sub> for HEK293 *versus* INS-1 832/3. “n.c.” indicates not calculable.

## Supplementary information

**Supplementary Figure 1.**
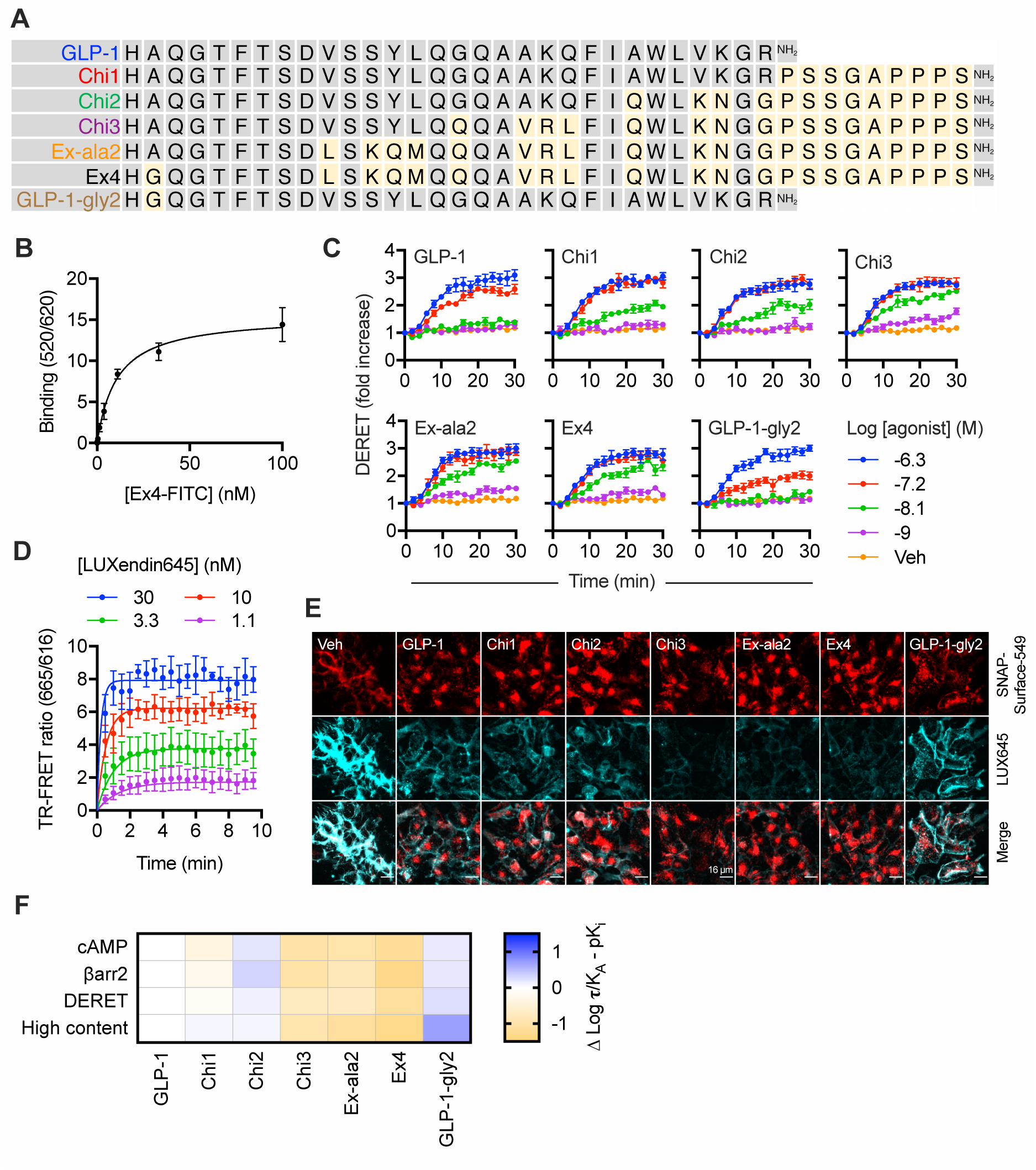
Additional binding, signalling, and trafficking data for chimeric GLP-1R ligands. (**A**) Peptide agonist sequences in single letter amino acid code, with exendin-4-specific residues highlighted in gold. (**B**) Saturation binding of exendin-4-FITC in HEK293-SNAP-GLP-1R cells, *n*=5, see also Figure 1B. (**C**) Kinetic traces for GLP-1R internalisation in HEK293-SNAP-GLP-1R cells stimulated with indicated concentration of agonist, measured by DERET, *n*=4, relates to Figure 1E. (**D**) TR-FRET measurements of LUXendin645 binding to Lumi4-Tb-labelled SNAP-GLP-1R in HEK293 cells, *n*=3, kinetic binding curve fitting of pooled data shown. (**E**) Widefield microscopy images of HEK293-SNAP-GLP-1R cells labelled with SNAP-Surface-549 prior to stimulation with 1 µM agonist for 30 minutes, followed by 60 minute recycling and labelling of surface GLP-1Rs with LUXendin645 (100 nM); representative images of *n*=5 independent experiments shown, with identical brightness and contrast settings across all single-channel images; scale bar = 16 µm. (**F**) Heatmap representation of coupling between occupancy and cAMP, β-arrestin-2 recruitment and endocytosis (measured by DERET and high content microscopy) signalling, determined by subtraction of pK_i_ from log **τ**/K_A_ values for each pathway (see Tables 2 and 3) and subsequently normalised to GLP-1. Data represented as mean ± SEM.

**Supplementary Figure 2.**
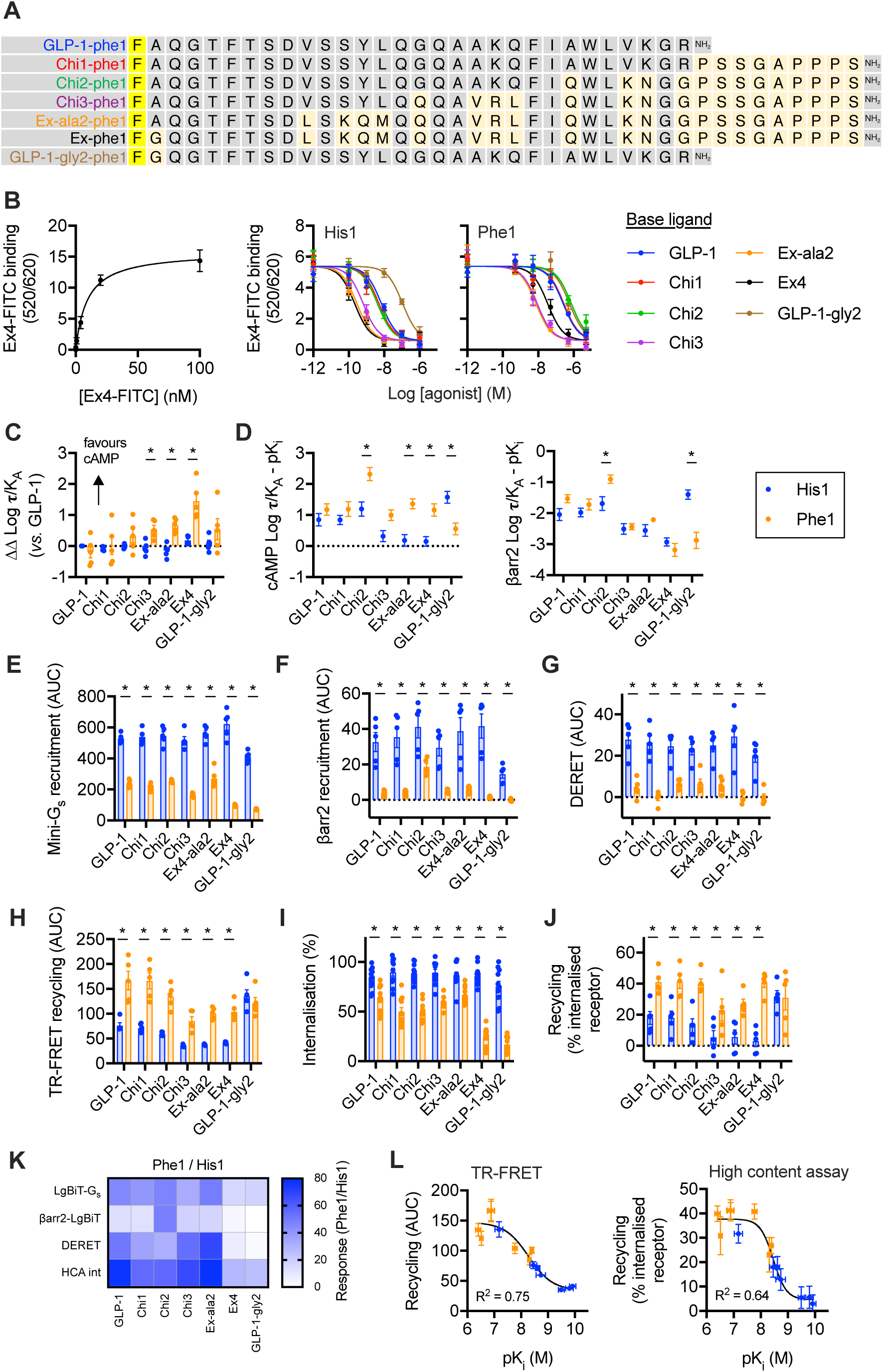
Phe1-substituted ligand evaluation. (**A**) Phe1 peptide sequences in single letter amino acid code, with exendin-4-specific residues highlighted in gold. (**B**) Equilibrium binding studies in HEK293-SNAP-GLP-1R cells, showing saturation binding of exendin-4-FITC measured by TR-FRET, with parallel measurements of 4 nM exendin-4-FITC binding in competition with indicated concentration of unlabelled agonist, *n*=5. (**C**) Alternative depiction of data shown by heatmap in Figure 2A, indicating bias (ΔΔ log **τ**/K_A_) of each ligand relative to GLP-1. (**D**) Representation of coupling between occupancy and cAMP and β-arrestin-2 responses in PathHunter CHO-K1-βarr2-EA-GLP-1R cells, determined by subtraction of pK_i_ from log **τ**/K_A_ values for each pathway (see Table 3), with error propagation, with each ligand pair compared by one-way ANOVA with Sidak’s test. (**E**) AUC analysis for LgBiT-mini-G_s_ recruitment (Figure 2B), with statistical comparison by one-way randomised block ANOVA with Sidak’s test to compare bias for each His1 / Phe1 ligand pair. (**F**) As for (E) but for βarr2-LgBiT recruitment (Figure 2C). (**G**) As for (E) but for internalisation measured by DERET (Figure 2D). (**H**) As for (E) but for GLP-1R recycling measured by TR-FRET (Figure 2E). (**I**) Quantification of GLP-1R internalisation in HEK293-SNAP-GLP-1R cells after 1 µM ligand treatment for 30 minutes, measured by high content microscopy analysis, *n*=11, with statistical comparison by one-way randomised block ANOVA with Sidak’s test to compare bias for each His1 / Phe1 ligand pair. (**J**) As for I, but for GLP-1R recycling after 1 µM ligand pre-treatment, *n*=5. (**K**) Alternative representation of data shown in Figures 2B, 2C, 2D and Supplementary Figure 2I, with Phe1 ligand responses expressed relative to the equivalent His1 ligand. (**L**) Relationship between agonist binding affinity (see Table 4) and GLP-1R recycling measuremed by TR-FRET (see Figure 2E) and high content microscopy (see Supplementary Figure 2J), with 4-parameter logistic fits and goodness-of-fit shown. *p<0.05 by statistical test indicated in the text. Data represented as mean ± SEM, with individual replicates shown in some cases.

**Supplementary Figure 3.**
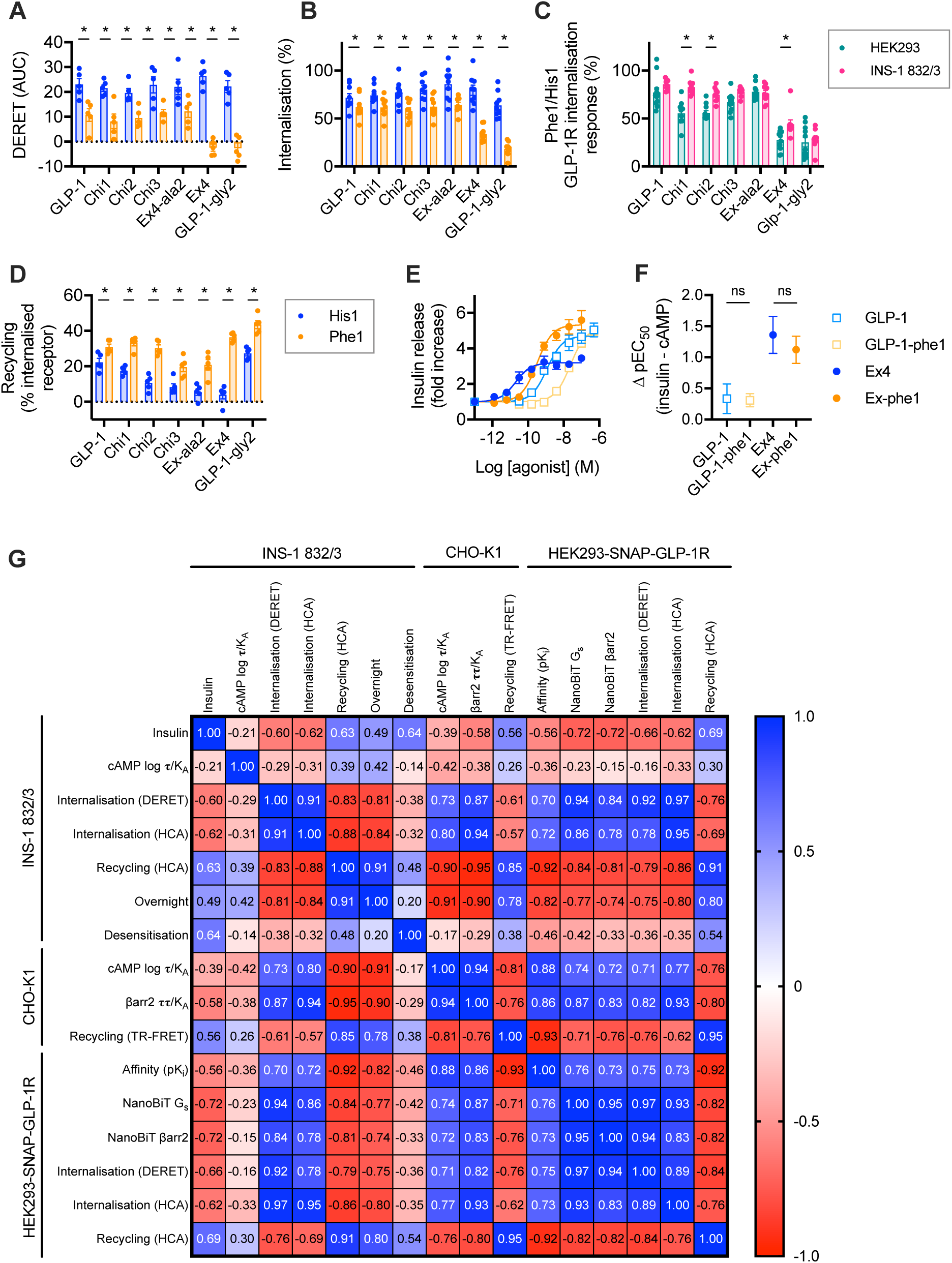
Effects in beta cells. (**A**) Alternative representation of heatmap data from Figure 3B, i.e. DERET-measured GLP-1R internalisation AUC in INS-1 832/3 GLP-1R^-/-^ cells stimulated with 1 µM agonist, *n*=5, statistically compared by one-way randomised block ANOVA with Sidak’s test for each His1 *versus* Phe1 ligand pair. (**B**) Quantification of SNAP-GLP-1R internalisation in INS-1 832/3 GLP-1R^-/-^ cells after 1 µM ligand treatment for 30 minutes, measured by high content microscopy analysis, *n*=9, with statistical comparison by one-way randomised block ANOVA with Sidak’s test to compare bias for each His1 / Phe1 ligand pair. (**C**) Comparison of Phe1 ligand internalisation measurements in INS-1 832/3 cells in comparison to HEK293 cells (see Supplementary Figure 2I), with the response of each Phe1 ligand expressed relative to that of its His1 counterpart for each assay, with statistical comparisons performed by one way ANOVA with Sidak’s test. (**D**) As for (B), but for GLP-1R recycling after 1 µM ligand pre-treatment, *n*=5. (**E**) Insulin secretion from wild-type INS-1 832/3 cells treated with 11 mM glucose ± indicated agonist dose for 16 hours, expressed relative to vehicle, *n*=5. (**F**) Comparison of potency estimates for acute cAMP signalling (Figure 3A, Table 5) and sustained insulin secretion (Supplementary Figure 3E) performed by subtraction pEC_50_ values, with error propagation, one-way ANOVA with Sidak’s test for each His1 *versus* Phe1 ligand pair. (**G**) Correlation matrix summarising relationship between agonist responses included in this work; single-dose responses are normalised on a 0 – 100% scale, whereas logarithmically quantified indices (pK_i_, log **τ**/K_A_) have not been further normalised; Pearson r coefficient is shown for each comparison. *p<0.05 by statistical test indicated in the text. Data represented as mean ± SEM, with individual replicates shown throughout.

## Acknowledgements

This work was funded by an MRC project grant to B.J., A.T., S.R.B. and G.A.R. The Section of Endocrinology and Investigative Medicine is funded by grants from the MRC, BBSRC, NIHR, an Integrative Mammalian Biology (IMB) Capacity Building Award, an FP7-HEALTH-2009-241592 EuroCHIP grant and is supported by the NIHR Biomedical Research Centre Funding Scheme. The views expressed are those of the author(s) and not necessarily those of the funder. B.J. was also supported by the Academy of Medical Sciences, Society for Endocrinology and an EPSRC capital award. D.J.H. was supported by MRC (MR/N00275X/1 and MR/S025618/1) and Diabetes UK (17/0005681) Project Grants. This project has received funding from the European Research Council (ERC) under the European Union’s Horizon 2020 research and innovation programme (Starting Grant 715884 to D.J.H.). G.A.R. was supported by a Wellcome Trust Investigator Award (212625/Z/18/Z), MRC Programme grants (MR/R022259/1, MR/J0003042/1, MR/L020149/1) Experimental Challenge Grant (DIVA, MR/L02036X/1), MRC (MR/N00275X/1), and Diabetes UK (BDA/11/0004210,

BDA/15/0005275, BDA 16/0005485) grants. This project has received funding from the European Union’s Horizon 2020 research and innovation programme via the Innovative Medicines Initiative 2 Joint Undertaking under grant agreement No 115881 (RHAPSODY) to G.A.R.

## Author contributions

Z.F., S.C., P.P, B.J. and A.T. and designed the study and performed experiments. J.B., D.J.H. and I.R.C. provided novel reagents. S.K., F.G., C.D. and P.F. developed software for high content image analysis. B.J., A.T., S.R.B., G.A.R. and T.T. acquired core funding for this project. B.J. and A.T. wrote the manuscript. All authors reviewed and approved the manuscript.

